# Highly multiplexed *in situ* protein imaging with signal amplification by Immuno-SABER

**DOI:** 10.1101/507566

**Authors:** Sinem K. Saka, Yu Wang, Jocelyn Y. Kishi, Allen Zhu, Yitian Zeng, Wenxin Xie, Koray Kirli, Clarence Yapp, Marcelo Cicconet, Brian J. Beliveau, Sylvain W. Lapan, Siyuan Yin, Millicent Lin, Edward S. Boyden, Pascal S. Kaeser, German Pihan, George M. Church, Peng Yin

## Abstract

Probing the molecular organization of tissues requires *in situ* analysis by microscopy. However current limitations in multiplexing, sensitivity, and throughput collectively constitute a major barrier for comprehensive single-cell profiling of proteins. Here, we report Immunostaining with Signal Amplification By Exchange Reaction (Immuno-SABER), a rapid, highly multiplexed signal amplification method that simultaneously tackles these key challenges. Immuno-SABER utilizes DNA-barcoded antibodies and provides a method for highly multiplexed signal amplification via modular orthogonal DNA concatemers generated by Primer Exchange Reaction. This approach offers the capability to preprogram and control the amplification level independently for multiple targets without *in situ* enzymatic reactions, and the intrinsic scalability to rapidly amplify and image a large number of protein targets. We validated our approach in diverse sample types including cultured cells, cryosections, FFPE sections, and whole mount tissues. We demonstrated independently tunable 5-180-fold amplification for multiple targets, covering the full signal range conventionally achieved by secondary antibodies to tyramide signal amplification, as well as simultaneous signal amplification for 10 different proteins using standard equipment and workflow. We further combined Immuno-SABER with Expansion Microscopy to enable rapid and highly multiplexed super-resolution tissue imaging. Overall, Immuno-SABER presents an effective and accessible platform for rapid, multiplexed imaging of proteins across scales with high sensitivity.

## Introduction

From C. *elegans* to humans, multicellular organisms are highly organized systems with large numbers of different cell types interacting to carry out biological functions. *In situ* protein imaging using immunofluorescence (IF) methods enables researchers to label specific cell types with affinity probes (e.g. antibodies) and visualize cells within their biological environment. For a comprehensive understanding of function and dysfunction, it is necessary to obtain spatial information of multiple markers that can accurately represent the type and state of individual cells, making highly multiplexed imaging a valuable capability. However, conventional IF methods offer limited multiplexing (typically of < 5 targets) due to spectral overlap of fluorophores.

Different methods have been developed to address this limitation, including multiplexed detection via unconventional probes and specialized instruments (as applied in multiplexed ion beam imaging [MIBI]^1, 2^, imaging mass cytometry [IMC]^3, 4^, or multiplexed vibrational imaging^5^), iterative sequential antibody labeling and imaging (MxIF^6^ and CycIF^7, 8^), and sequential detection through DNA-barcoding (DNA-Exchange-Imaging [DEI]^9^ and CO-Detection by indEXing [CODEX]^10^). These techniques can achieve higher multiplexing (i.e. ≥5-targets), but typically at an expense of decreased sensitivity, throughput, or accessibility (specialized instruments). MIBI, IMC and vibrational imaging requires specialized instruments and typically operate by point-scanning small fields of view (∼2-5 min per 50 µm × 50 µm field of view); hence are exceedingly slow for large areas like centimeter scale tissue sections^1, 5^. In contrast, fluorescence methods utilizing sequential antibody staining (MxIF and CycIF) offer high accessibility and relatively fast image acquisition; however, they require multiple cycles of slow primary antibody incubation steps (typically hours to overnight per staining cycle), and can result in total experimental time of weeks to months (scaling with the number of targets)^6-8^. As a recent alternative, the development of DNA-barcoding based methods has created an important advance in throughput, while preserving high accessibility. In this case, multiplexing is enabled by the orthogonality of pre-designed DNA sequences, allowing ten(s) of antibodies to be applied simultaneously, followed by fast sequential readout of the sequences either through rapid binding/unbinding of short fluorescent oligonucleotides using DNA-Exchange technique^9, 11-14^ or by fluorescent dNTP analogs and *in situ* polymerization as in CODEX^10^.

DNA-barcoding of antibodies as in DEI or CODEX, allows staining total staining/multiplexed imaging time of any sample size can be reduced to a couple of days, however, to overcome the overlap of antibody host species, high multiplexing requires direct conjugation of primary antibodies with DNA oligonucleotides. This causes signal levels to be lower due to the lack of signal amplification from polyclonal secondary antibodies and use of single fluorophore labeled probes. Low signal level, in turn, hampers the detection sensitivity, and makes it hard to visualize low abundance targets especially in tissues with high autofluorescence and scattering, and low antigen access. Limited signal also imposes longer acquisition times (due to lengthy exposure times or slow scanning speeds), and lowers the throughput of imaging (and/or resolution due to the need to integrate the signal over several pixels). Incorporating *in situ* signal amplification is hence critically needed to generate higher signal for each target and support higher throughput and sensitivity.

For RNA molecules, amplification with high-multiplexing has been attained via combinatorial barcoding or amplicon-supported *in situ* sequencing^15-18^. However, these capabilities have not translated similarly for multiplexed amplification of protein targets, owing to differences in their cellular organization. Unlike RNA transcripts, which are sparsely organized and mostly found as discrete spots, proteins are spatially overlapping and are organized in clusters with high densities of multiple other targets. This makes the combinatorial or sequencing based readout of barcodes in multiplexed fashion very challenging. Additionally, the dynamic range of the cellular proteome approaches seven orders of magnitude, from a few copies to ten million copies per cell, yielding a range 1000-fold higher than that observed for the transcriptome^19-22^. Hence it is desirable that signal amplification for proteins would be not only (i) compatible with fast multiplexing and easy to scale-up, but also (ii) applicable for spatially overlapping, high density organizations of the targets, and (iii) simultaneously tunable in a user defined manner for each target. Currently available *in situ* signal amplification methods pose limitations for satisfying these requirements.

Tyramide signal amplification (TSA)^23, 24^ provides an effective way for amplification of the signal through covalent binding of the diffusive tyramide molecules in the vicinity of the target. But, due to lack of orthogonal chemistries, TSA is restricted to labeling of one target at a time. Labeling multiple targets requires sequential antibody labeling and signal amplification with different fluorophores need to be performed sequentially with microwave-based removal of antibodies after each round (up to 7-color multispectral imaging has been demonstrated this way)^25-27^. Relying on the orthogonal sequence design, DNA-mediated methods offer a promising route for simultaneous signal amplification. DNA-tagged antibodies allows for signal amplification through either DNA template copying or component self-assembly. In rolling circle amplification (RCA)^28^, a processive polymerase acts on a circular template to synthesize long concatenated repeats, generating micron-diameter balls of DNA. RCA offers high levels of amplification and potential for multiplexing, but *in situ* enzymatic reaction is hard to control or tunable for individual targets^29^. Both TSA and RCA may also lead to blurring of signals and decreased resolution, respectively due to spreading of the tyramide molecules or the large size of the amplicons (reaching from 250 nm to over ∼1 µm radius^30, 31^). Branched DNA assemblies^32-34^, such as RNAscope^35, 36^, generate complex tree structures for stable binding of fluorescent DNA strands, whereas HCR utilizes the triggered assembly of metastable fluorophore-conjugated hairpins^37-39^. The structural complexities of these existing DNA-assembly based platforms present potential challenges for designing highly multiplexed orthogonal systems (e.g. each HCR design entails working with the complex kinetic pathway of triggered assembly of two meta-stable hairpins^40-42^). In practice, simultaneous signal amplification for proteins beyond spectral multiplexing (3-5 targets) remain to be demonstrated^43, 44^.

Therefore, a robust method that simultaneously allows highly multiplexed and individually controllable signal amplification for sensitive *in situ* protein detection is of pressing need. Here we report a highly multiplexed signal amplification method that we name **<u>Immuno</u>staining with <u>S</u>ignal <u>A</u>mplification <u>b</u>y <u>E</u>xchange <u>R</u>eaction (Immuno-SABER)** (**Fig. 1**). In order to amplify the signal of DNA-barcoded antibodies, we utilize single-stranded DNA concatemers that are generated in a pre-programmed manner via our recently engineered Primer Exchange Reaction (PER)^45^ **(Fig. 1a).** These concatemers provide controllable multiplexed signal amplification by acting as docking sites for multiple fluorophore-bearing DNA imager strands^46^ **(Fig. 1b)**. We achieve rapid multiplexing through single-step immunostaining with DNA-barcoded primary antibodies followed by sequential hybridization and dehybridization of DNA imager strands (**Fig. 1c**). Single step staining of all DNA-conjugated primary antibodies eliminates the need of secondary antibody species pairing and repeated antibody retrieval and staining. Robust signal amplification enables fast imaging by allowing reduced image acquisition times.

**Figure 1.**
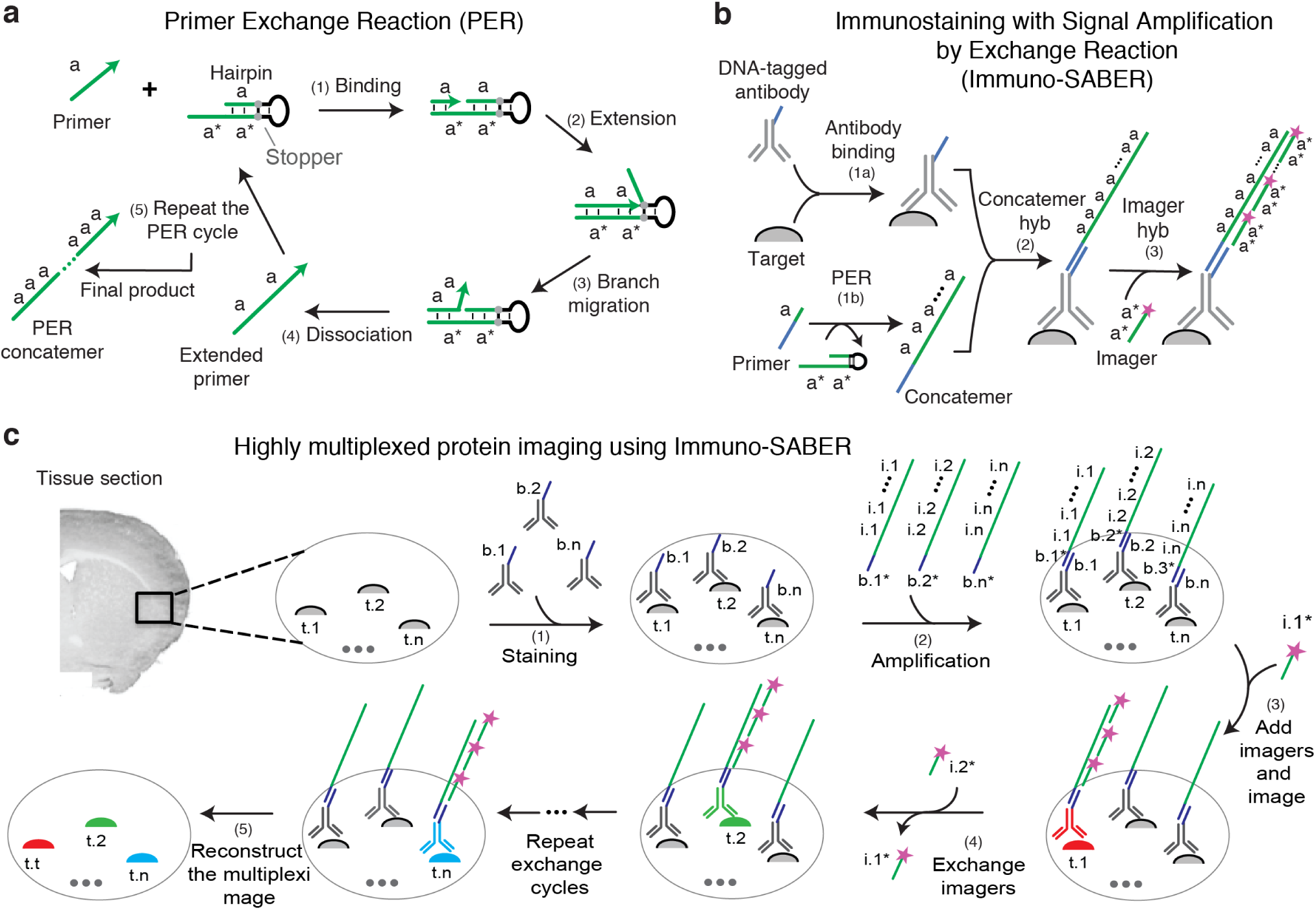
Immuno-SABER scheme. **(a)** Mechanism of PER^45^: (1) A 9-mer primer of sequence *a* binds to the single stranded *a*\* sequence on the hairpin. Asterisk denotes complementarity. (2) The primer is extended by a strand displacing polymerase (e.g. Bst) in an isothermal and autonomous manner. Each PER hairpin features a stopper sequence that halts polymerization and results in release of the polymerase. (3) The newly synthesized *a* segment is displaced from binding with the *a*\* segment of the hairpin through branch migration. (4) The extended primer and the intact hairpin autonomously dissociate. (5) Repetition of this copy-and-release process through repetitive reaction cycles produces a long concatemer of *a*. This simple step-by-step synthesis of concatemers offers a tight control of the reaction by external parameters (such as hairpin or dNTP concentration, reaction time and temperature) as well as high programmability. **(b)** Schematic of Immuno-SABER: (1a) Antibodies are conjugated with DNA bridge strands (sequences) and used to simultaneously stain multiple targets in biological samples as in DEI^9^. (1b) Primer sequences (green line) are extended to a controlled length using PER. (2) The concatemers are hybridized to the bridge sequence (blue) on the antibody. (3) Fluorophore (depicted as purple star)-labeled 20-mer DNA “imagers” strands hybridize to the repeated binding sites on the long PER concatemers. Each imager is designed to bind to a dimer of the unit primer sequence. **(c)** Schematic of highly multiplexed imaging using Immuno-SABER: (1) Different biological targets (t.1 to t.n) are labeled with corresponding antibodies conjugated with orthogonal DNA bridge strands (b.1 to b.n). (2) Orthogonal pre-extended concatemers are hybridized (via bridge complements b.1* to b.n*) to create simultaneous signal amplification. (3) The targets are visualized by hybridization of fluorophore-labeled DNA imaging strands (i.1* to i.n*) to their repeated binding sites (i.1 to i.n) on the orthogonal concatemers. (4) Multiple targets can be imaged by rapid exchange rounds, where orthogonal imagers are dehybridized and hybridized in multiple cycles. (5) The images are computationally aligned and pseudo-colorized to overlay different targets in the same sample.

We validate Immuno-SABER in diverse sample types including cultured cells, thick cryosections, thin FFPE sections and whole mount tissue samples, and present the capability to preprogram and control the amplification level independently for multiple targets and without *in situ* enzymatic reactions. Achieving independently tunable 5-180-fold amplification, Immuno-SABER unlocks the full signal range conventionally achieved by secondary antibodies up to tyramide signal amplification. The simple design criteria of PER sequences make Immuno-SABER intrinsically scalable. In practice, we performed an *in situ* orthogonality assay for a subset of our previously designed library of 50 PER sequences^46^, and observed crosstalk for only one pair. We further demonstrate simultaneous signal amplification and imaging of 10 protein targets in tissues. Finally, we combine SABER with Expansion Microscopy (ExM)^47^ to achieve rapid, highly multiplexed super-resolution imaging for tissues.

In addition to its unique performance for highly multiplexed signal amplification for protein targets in an individually controllable fashion, Immuno-SABER further features a simple workflow, utilizes low cost reagents, and is compatible with existing standard imaging platforms in common labs. Thus, Immuno-SABER presents an effective and accessible platform for *in situ* spatial mapping of proteins in tissues across scales. By putting DNA-mediated highly multiplexed amplification into practice, Immuno-SABER offers an important stepping-stone for realizing the promise of DNA-mediated imaging. Together with our complementary recent work^46^, here we generate novel concepts for pre-programmed *in vitro* synthesis and directed *in situ* assembly that enables highly multiplexed amplification in an accessible, low-cost and open-source manner leading the way for scaling up the capabilities of DNA-mediated imaging. These concepts can be adopted by existing DNA-based methodologies of labeling and amplification to yield a plethora of future methods and applications.

## Results

### Validation and quantification of in situ signal amplification using Immuno-SABER

For *in situ* signal amplification, Immuno-SABER relies on a strategy entailing controlled *in vitro* synthesis of amplifier concatemers by PER, followed by programmed *in situ* assembly. PER utilizes a catalytic hairpin for controllable extension of a short primer sequence in an iterative manner (as depicted in **Fig. 1a**). Using PER, we synthesized long DNA concatemers of desired lengths reaching >500 bases through modulation of reaction conditions such as reaction time, hairpin or dNTP concentration^46^. We first tested the suitability of these long DNA concatemers for *in situ* imaging in biological samples and quantified the amplification capability. To have a modular design for protein imaging, we utilized 42-nucleotide bridge sequences that would enable coupling of concatemers of choice to antibodies on demand. We provided the design criteria to create orthogonal bridge sequence libraries for probe barcoding previously^46^. We conjugated these 42mer single-stranded DNA (ssDNA) oligos to antibodies targeting lysine residues (see Methods for details). We also developed an optional DNA-tag and toehold displacement mediated purification strategy **(Fig. S1)**. The PER primers we utilize are similarly barcoded with complements of these bridge sequences. These primers are extended into concatemers via PER *in vitro*, and then applied onto cells and hybridized to the bridges *in situ*. For application of this *in vitro* extension and *in situ* assembly strategy on biological specimens there main considerations are: i) specificity of the labeling, ii) ability to label dense targets (due to potential interference of the probe size), iii) penetration and access to targets in highly crosslinked or thick tissue samples, and iv) efficient amplification for diverse targets. To validate our approach and mitigate these potential considerations we have performed several demonstrations, and in each case included controls for a case where the same antibody was used without amplification or a conventional staining was performed with fluorophore-conjugated secondary antibodies **(Fig. 2a)**. First, to evaluate the specificity and preservation of morphology upon labeling with the linear concatemers, we performed Immuno-SABER staining in cultured cells for microtubules as a test case for a densely arranged structural protein target. We have observed specific staining, clear tubular morphology and similar staining pattern to conventional immunostaining with fluorophore-conjugated secondary antibodies **(Fig. 2b)**. The resolution of the images, as quantified by full-width-half-maximum (FWHM) of the fitted line plots across the thin microtubules, was unaltered compared to the secondary antibody control **(Fig. S2a-c)**. These results demonstrated the suitability of Immuno-SABER for *in situ* labeling of proteins, with high resolution and high specificity even at dense arrangements.

**Figure 2.**
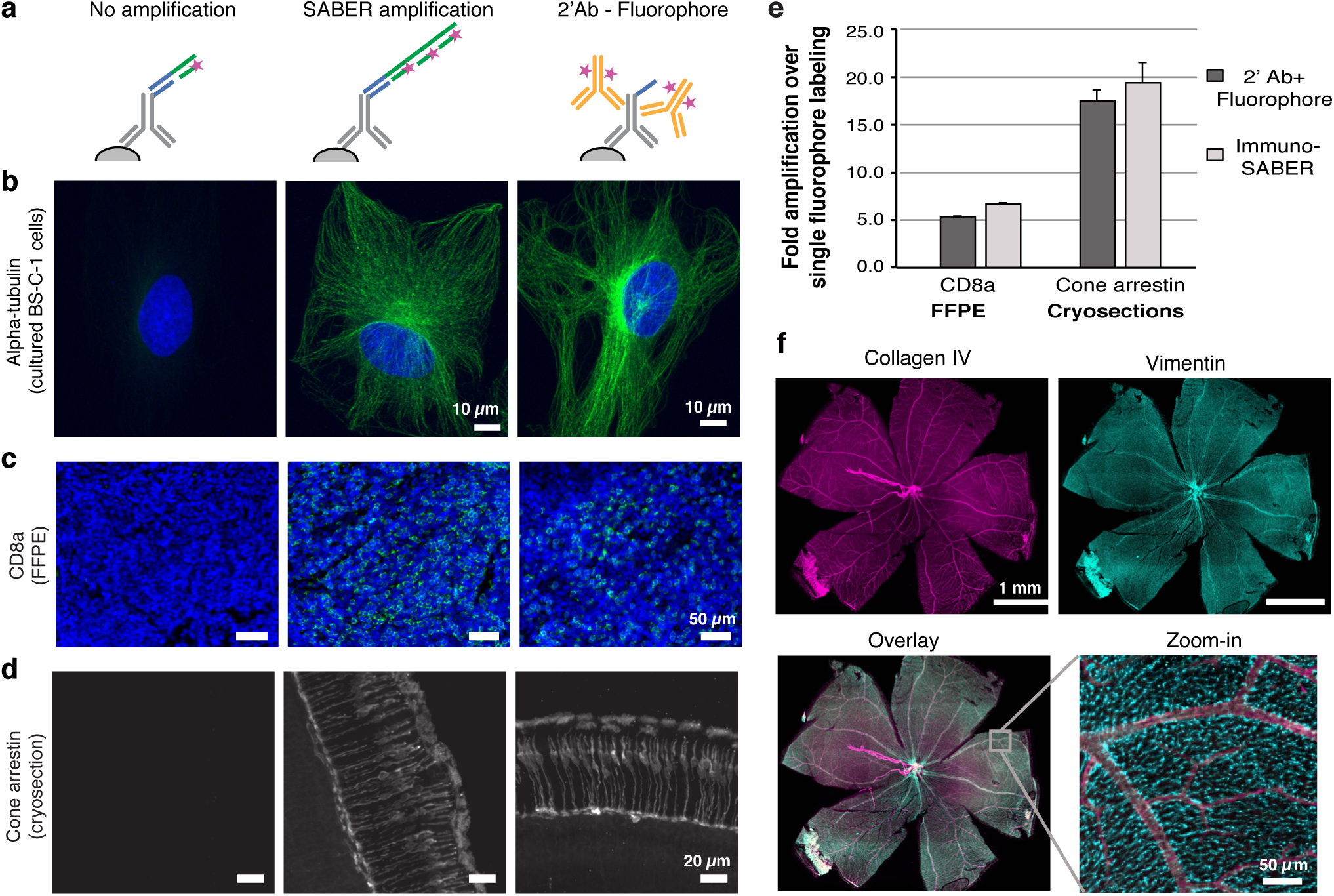
Validation and quantification of signal amplification by Immuno-SABER. **(a)** Cultured BS-C-1 cells were immunostained for alpha-tubulin and three conditions were prepared for comparison: Unamplified condition, where (i) unextended primers with single binding site for imager (with Alexa647) was hybridized to the bridge on the antibody, (ii) the extended concatemer was hybridized for signal amplification (linear amplification), (iii) conventional antibody staining was performed with Alexa647-conjugated secondary antibodies. **(b)** Representative images of each (max projections from confocal z-stack taken with a 63× objective). **(c)** Representative images for CD8a staining (labeled with ATTO488-imager) in human tonsil FFPE sections (single plane large area scans with a 20× objective cropped to show a region of the CD8a^+^-cell rich interfollicular zone. **(d)** Cone arrestin staining in mouse retina cryosections (maximum projection of a confocal z-stack taken with a 20× objective). See Methods for experimental details. **(e)** Level of signal amplification by Immuno-SABER was quantified by measuring the background-subtracted mean fluorescence for several regions of interest in the tissues and expressed as fold amplification over unamplified signal level. Conventional secondary antibody amplification was also quantified similarly and shown as reference. For CD8a, n = 90-93 rectangular ROIs (each covering 0.03-1.20 mm^2^ tissue regions; consecutive sections are used for the three conditions). For cone arrestin, n = 6 images. Error bar indicates SEM. **(f)** Immuno-SABER was performed in whole-mount retina sections for collagen IV and vimentin. Maximum projections from confocal z-stacks are displayed.

For validation of the labeling strategy in tissue samples and quantification of the signal amplification capability of Immuno-SABER, we used two types of tissue preparations: 5 µm-thick FFPE human tonsil sections and 40 µm-thick mouse retina cryosections. Both tissue types provide a good proof-of-principle system to develop a multiplexed imaging methodology thanks to morphologically distinct and well-conserved organization of different cell types that can be identified by well-established biomarkers. Retina has a layered organization of different types of cells whereas tonsils feature multiple germinal centers with precise stereotypic organization of a large number of distinct cell types.

For the FFPE samples, we imaged the T-cell membrane marker CD8a. CD8a^+^ cells largely are known to be present in the marginal zone (outside of the germinal centers)^48^. We were able to reproduce this organization with Immuno-SABER staining **(Fig. 2c)**. For mouse retina cryosections, we imaged cone arrestin, which is a specific marker of the cone photoreceptor cells^9^. Both markers were successfully imaged using Immuno-SABER, and the signal patterns were consistent with the expected distribution of the markers, suggesting high specificity of our method **(Fig. 2d)**. For both targets and sample types, Immuno-SABER yielded similar or slightly higher fluorescence signal than conventional fluorophore-conjugated secondary antibody staining using the same fluorophore. We quantified the signal amplification level using Immuno-SABER as 6.7-fold for CD8a and 19.4-fold for cone arrestin (see Methods for details of the quantification for the two sample types). For comparison, conventional secondary antibody staining yielded 5.3 and 17.5-fold, respectively **(Fig. 2e)**. Overall, our strategy yielded ∼5-20-fold improvement in the signal level compared to the unamplified control, yielding amplification at a similar level provided by secondary antibodies. We note that the degree of *in situ* signal amplification depends on multiple factors, including abundance and organization of targets, the antibodies (e.g. clonality, conjugation efficiency), the method of quantification (the unamplified signal level, thresholding, background subtraction), as well as the experimental conditions and properties of the SABER sequences (e.g. the length of SABER concatemer^46^).

Despite the anticipated size of the long concatemers reaching >500 nucleotides, SABER concatemers can effectively penetrate relatively thick samples. We validated the penetration capability of the concatemers in whole-mount preparations of mouse retina by successfully staining for the Muller cell marker Vimentin and blood vessel marker Collagen IV, which both predominantly localize in the 100 µm region from nerve fiber layer to outer plexiform layer of the retina, as expected^49^ **(Fig. 2e and S2d)**. This high penetration may potentially be attributed to SABER concatemers being largely linear DNA structures designed in 3-letter code (made of only A, T, and C nucleotides) to be devoid of secondary structures (that may be otherwise stabilized by G-C pairings).

### Enhancement of signal through branching

Although the amplification obtained through Immuno-SABER is substantial, further enhancement of the signal can be desirable for proteins of lower abundance, or to further improve the throughput of imaging (by allowing shorter exposure times through signal amplification). For this purpose, we developed a sequential amplification strategy where independently extended secondary concatemers can be branched off the primary concatemer to create more binding sites for fluorescent imagers (termed as ‘branched SABER’). This can be achieved by a sequential round of concatemer hybridization as depicted in **Fig. 3a**. We have performed similar tests to check the effect of an additional amplification round targeting the same proteins in cell culture, human FFPE tonsil tissues and mouse retina cryosections **(Fig. 3b-d and S3a-b**). Microtubule staining in cultured cells indicates that the specificity of labeling and the high-resolution morphology of the target were still preserved with branched SABER amplification **(Fig. S3a**). FWHM estimations for microtubule samples demonstrated that the size of the structures was still bound by the diffraction-limit of the imaging setup and were not substantially altered **(Fig. S3b**). With one round of branching, we obtained additional 2.8-fold amplification for CD8a in FFPE sections and 8.4-fold amplification for cone arrestin in cryosections over single round Immuno-SABER detection **(Fig. 3d**).

**Figure 3.**
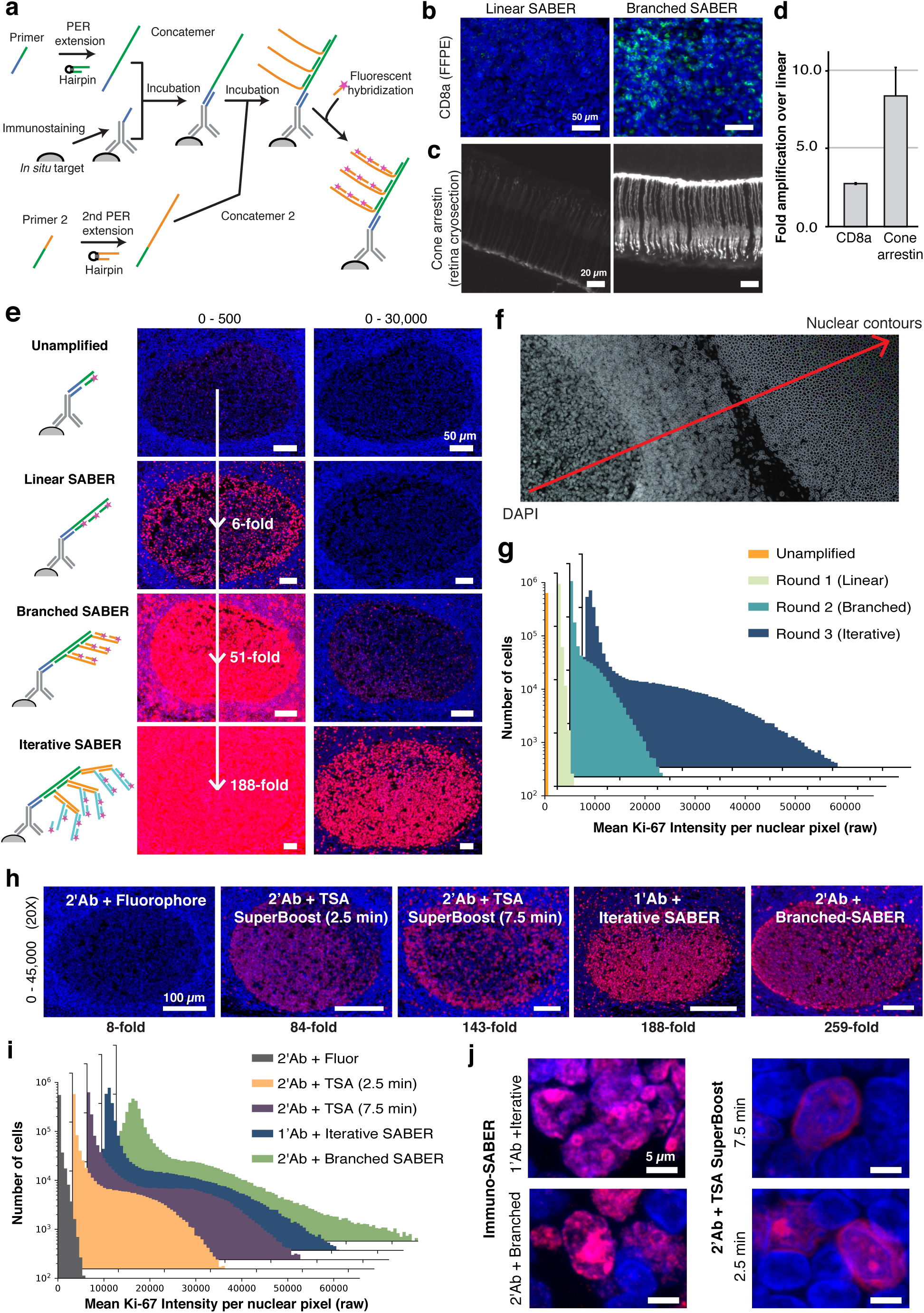
Immuno-SABER signal can be further amplified through branching. **(a)** Primary PER concatemers can be targeted by secondary concatemers to form a branched structure which amplifies the signal further by presenting additional binding sites for the imagers. **(b-c)** Representative images for linear and branched SABER amplification are shown for different preparations: **(b)** CD8a staining in human tonsil FFPE sections (single plane large area scans with 20× objective, cropped to show a CD8a-rich interfollicular zone region, **(c)** Cone arrestin staining in mouse retina cryosections (max projections from confocal z-stacks). **(d)** Level of signal amplification by branched Immuno-SABER over linear was quantified by measuring the background-subtracted mean fluorescence for several regions of interest in the tissues and expressed as fold amplification over linear SABER. For CD8a, n = 84-144 rectangular ROIs (each covering 0.03-1.20 mm^2^ tissue regions; consecutive sections are used for the two conditions). For cone arrestin, n = 6 images. Error bars are SEM. **(e)** Representative images for nuclear Ki-67 (Alexa647, red) with DAPI (blue) staining cropped to show a Ki-67 rich germinal center in FFPE human tonsil sections, with up to 3 rounds of amplification (iterative SABER). 16-bit images were scaled to two different maximum pixel values (500 and 30,000) to allow visual comparison. Quantification of the signal amplification in each case versus the unamplified sample was performed at the whole tissue section level, by segmenting individual nuclei and using the total nuclear intensity for each sample (without background subtraction), as described in panel **g-h** and Methods. Fold amplification values are provided with the images. **(f)** Machine-learning based highlighting of nuclear contours from DAPI staining images. The deep learning model was trained with a manually annotated dataset to enable automatic identification of nuclear contours to be followed by watershed segmentation **(Fig. S3g)**. **(g)** Mean Ki-67 signal intensity for each nucleus was obtained from automated segmentation algorithm and the histogram was plotted for the whole tissue section for each condition. The sections each contain 636,479-717,176 cells. **(h)** Images show germinal centers in FFPE human tonsil sections with Ki-67 labeling (red) with conventional secondary antibody-fluorophore staining, with TSA, with Immuno-SABER using secondary antibodies with branching, or primary Ki-67 antibody with iterative SABER amplification (as in panel **e**). TSA was applied using poly-HRP conjugated secondary antibodies of Thermo SuperBoost Kit with 2.5 or 7.5 min tyramide-Alexa 647 incubation. The amplification level is noted below the images (as in **e**). **(i)** Histograms to visualize mean nuclear signal level were plotted (as in **g**) for the conditions in panel **i**. The consecutive tissue sections each contain 586,183-717,176 cells. The data displayed is part of the dataset in **e**. **(j)** Samples were imaged with a confocal microscope at higher resolution with 63× magnification to evaluate signal blurring. Images with different scaling are displayed in **Fig. S3h-i**.

Immuno-SABER allows further improvement of sensitivity by performing multiple hybridization rounds (termed as Iterative SABER). We applied this strategy both in tonsil FFPE sections and mouse retina cryosections, and performed 3 sequential rounds of SABER amplification following primary antibody staining. We targeted the proliferating cell nuclear marker Ki-67 that is mainly found in the germinal center (i.e. the dark zone) in tonsil sections, and synaptic marker SV2 that is found mainly in synapses of both outer and inner plexiform layers^9^. **Fig. 3e** shows Ki-67-rich germinal centers stained with indicated methods but imaged under the same microscope setting, and displayed at two different contrast levels to allow visual comparison of the signal level at every cycle.

To follow how the signal level changes for individual nuclei in the whole tissue section scans, we developed a machine-learning algorithm that automatically annotates DAPI-defined nuclear contours **(Fig. 3f)** for segmentation of individual nuclei in these densely populated sample types. Nuclear segmentation allowed us to assign mean Ki-67 signal intensity to each individual nucleus from whole slide scans with 550,000-750,000 cells each. To visualize the change of signal level, histograms were plotted for each round of branching using SABER **(Fig. 3g)**. We also utilized the total nuclear signal level for each condition to obtain a basic quantitative estimate of the signal amplification with respect to the unamplified case, noted as fold-amplification values over the respective images in **Fig. 3e**.

For the branching experiments on 5 µm FFPE sections, performing the concatemer incubations for longer periods (overnight for the longer primary concatemer, and 3 h for the branching round as for **Fig. 3e**) did not amplify the signal better than shorter (75 minutes each) concatemer incubation rounds (**Fig. S3c-d**), suggesting faster iterative amplification could be performed to shorten the total experimental time. Iterative-SABER (3-round) was also tested in thicker SV2-stained mouse retina cryosections yielding ∼80-fold amplification over unamplified control **(Fig. S3e).**

For such high levels of amplification, catalyzed reporter deposition-based TSA is considered a gold-standard, which was reported to achieve 10-1000-fold sensitivity improvement under different assay types and comparison conditions^24, 50-53^. This can be further improved 2-10 fold by using poly-HRP conjugated antibodies. However, due to lack of orthogonal chemistries TSA can only be applied to one target at a time, so to label multiple targets sequential antibody labeling and signal amplification with different fluorophores need to be performed with microwave-based removal of antibodies after each round^25-27^. In addition, the amplification level in TSA is difficult to control and it is not ideal for high-resolution imaging due to spreading of the fluorescent tyramide molecules to nearby areas (reaction products have been shown to spread over a ∼1 µm radius^30^). To investigate how Immuno-SABER performs in comparison to TSA, we first utilized the conventional TSA Kit from ThermoFisher that used mono-HRP conjugated secondary antibodies and -Alexa647-tyramide (i.e. signals were amplified both by secondary antibodies and by TSA). Although only DNA-conjugated primary antibodies were used, we found that iterative SABER could amplify signals to a higher level than TSA with secondaries (**Fig. S3f)**. For a more thorough comparison, we also used the SuperBoost kit from ThermoFisher that utilized poly-HRP conjugated secondaries for Ki-67 staining of FFPE tonsil sections, and compared the signal level obtained with iterative SABER (3 rounds of amplification) with primary antibodies, as well as branched SABER (2-rounds of amplification) using secondary antibodies, where we respectively achieved 188 and 259-fold amplification over the unamplified condition **(Fig. 3h)**. Both cases yielded higher signal improvement than TSA with poly-HRP conjugated secondary antibodies (up to 143-fold, **Fig. 3h-i**). We also note that at this high amplification level, Immuno-SABER still provided crisp staining and preserved the subcellular morphology, whereas TSA-staining resulted in blurring of the signal and spilling over of the label from the target compartments (Ki-67 is a nuclear protein highly enriched in nucleoli), which can be visualized as Ki-67 signal coming from outside of DAPI labeled nuclei in the high-magnification confocal images of single cells **(Fig. 3j, Fig. S3h-i**). Increasing the tyramide incubation duration to 10 minutes (2-10 minutes is the suggested optimization range by the manufacturer) worsened the blurring (**Fig. S3h, S3i rightmost panel**).

### Validating sequences for multiplexing Immuno-SABER

PER has little inherent restrictions in sequence design outside of preferred single strandedness of the concatemer. We previously developed a computational pipeline to design orthogonal PER primer-hairpin pairs with maximum extension efficiency and minimum crosstalk by utilizing *in silico* simulations using NUPACK^54^ and designed 50 orthogonal sequences to enable multiplexed imaging^45, 46^ Here, we tested the performance to 32 SABER sequences to extend into long DNA concatemers in controlled fashion using an *in vitro* gel-shift assay. **Fig. 4a** displays long concatemers extended from 32 primers up to 600 to 700-nucleotide lengths (which is the range we predominantly use for primary SABER concatemers^46^), visualized with SybrGold staining on denaturing PAGE gels. Since extension efficiency is sequence-dependent, reaction conditions were optimized by modulating the hairpin concentration and reaction time for each primer to obtain concatemers of similar lengths. All sequences in our test library succeeded to produce concatemers in the desired length range. All of the tested primers extended into long concatemers, with 31 out of 32 sequences (except #51) yielding a predominant long concatemer band at the desired target length, albeit some heterogeneity in the distribution of shorter reaction products **(Fig. S4a)**.

**Figure 4.**
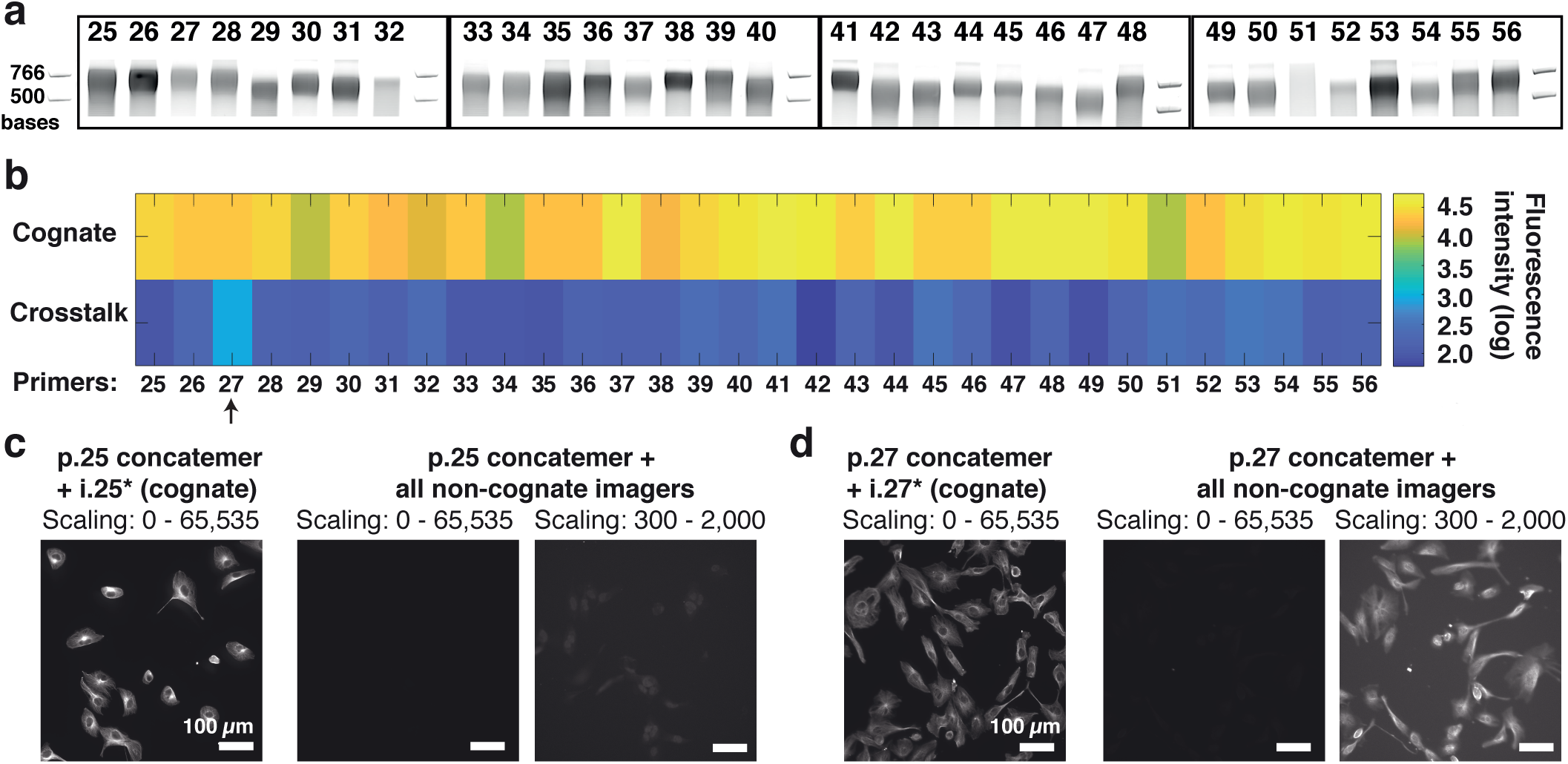
Sequence validation for highly multiplexed Immuno-SABER. **(a)** 32 SABER sequences were extended to ∼650 bases *in vitro* and examined by gel shift assay on a 6% denaturing PAGE gel. The 500 and 766 base ladder bands are displayed for reference. Full gel images are available in **Fig. S4a**. **(b)** *In situ* performance and crosstalk analysis of SABER sequences. BS-C-1 cells were stained with bridge DNA-conjugated antibodies targeting α-Tubulin on a 96-well plate. Concatemers extended from each primer were hybridized to the bridges creating an array of wells labeled with primer sequences p.25-p.56. For each primer (e.g. p.25), both cognate and crosstalk wells were prepared by either applying the corresponding Alexa647-imager (e.g. i.25* for the concatemer extended from p.25) or by applying all the imagers except the cognate one (e.g. -i.26* to i.56* for p.25 concatemer). See the Methods for details of the sequences. Images were captured in 16-bit (0-65,535). The fluorescence signals were quantified and plotted in the log scale and displayed as a heatmap. Non-negligible crosstalk signal was only detected for Primer 27 concatemer (indicated by the arrow). **(c-d)** Representative images are shown for Primer 25 (p.25) and Primer 27 (p.27) concatemers. Crosstalk images are displayed with two different intensity scales to render the crosstalk signal visible.

After *in silico* design and *in vitro* validation of SABER sequences, we evaluated the *in situ* performance and orthogonality of detection (crosstalk check) through an imaging-based multi-well plate assay (**Fig. 4b and c)**. DNA-conjugated antibodies targeting α-Tubulin were used to stain microtubules in fixed BS-C-1 cells, followed by hybridization of each DNA concatemer in separate wells. We divided the wells stained with the series of concatemers into two groups: (i) cognate group to be incubated with the corresponding imager strands, and (ii) crosstalk group where we added mixtures of imagers except the cognate imager strand. We were able to observe microtubule staining for all 32 sequences. Consistent with the *in vitro* gel shift assay, sequences that had lower extension efficiency (particularly primers 5, 8, 10 and 27) tended to yield less fluorescence signal compared to sequences that that extend with higher efficiency. We only observed minor crosstalk (crosstalk signal / cognate signal = 4%) for primer 27 (p.27). In the follow-up experiments, we determined the strand responsible for crosstalk as imager strand 48 (i.48*), which is excluded from the library for further multiplexed imaging (**Fig. S4b**).

### Multiplexed Immuno-SABER imaging in human tonsil FFPE sections

To validate the applicability of our method of applying independently programmed multiplexed signal amplification, we first tested spectral multiplexing on 5 µm human FFPE tonsil sections. FFPE samples are the standard preparations for clinical settings and for archival purposes. However, the preparation procedure results in increased autofluorescence and loss of antigenicity, which needs to be recovered (at least partially) through antigen retrieval protocols. Clinical immunohistochemistry procedures are typically applied for one target protein per section and utilize amplification methods such as chromatic reactions to improve the sensitivity. Multiple thin sections have to be prepared in order to stain multiple markers, which is not ideal since biopsy-based sample collection is invasive and samples can be scarce. Moreover, imaging throughput is a big concern for these samples since both for clinical pathology and for research studies the preference is to scan centimeter-scale tissues with whole slide scanning, potentially followed by higher resolution investigations of selected smaller regions of interest.

After standard preparation protocols for deparaffinization and antigen-retrieval, we simultaneously stained the samples for Ki-67, CD8a, IgA and IgM using antibodies conjugated to orthogonal bridge sequences. Among these targets IgA and IgM are expressed at high levels, whereas Ki-67 and CD8a are more moderately expressed (as a proxy, in the tonsil tissue RNA expression level of IgJ, the joining chain domain for multimeric IgA and IgM was found to be 1360.4 tags per million, while Ki-67 and CD8a expressions are at 15.1 and 18.9 tags per million, respectively^55, 56^). We performed whole slide scanning with 5-color spectral multiplexing (DAPI + 4 targets) **(Fig. 5a)**. In addition to multiplexing, signal amplification by Immuno-SABER allows improved imaging throughput by allowing short exposure times. For abundant proteins IgA and IgM, linear amplification was enough to get bright signal intensities that allow 1-20 ms exposure times. For moderately available proteins Ki-67 and CD8a branched SABER (two rounds of amplification) enabled 2-10 ms camera exposure times at 20× magnification. With these settings, we were able to obtain fast high-quality scans with subcellular resolution **(Fig. 5a, right)**.

**Figure 5.**
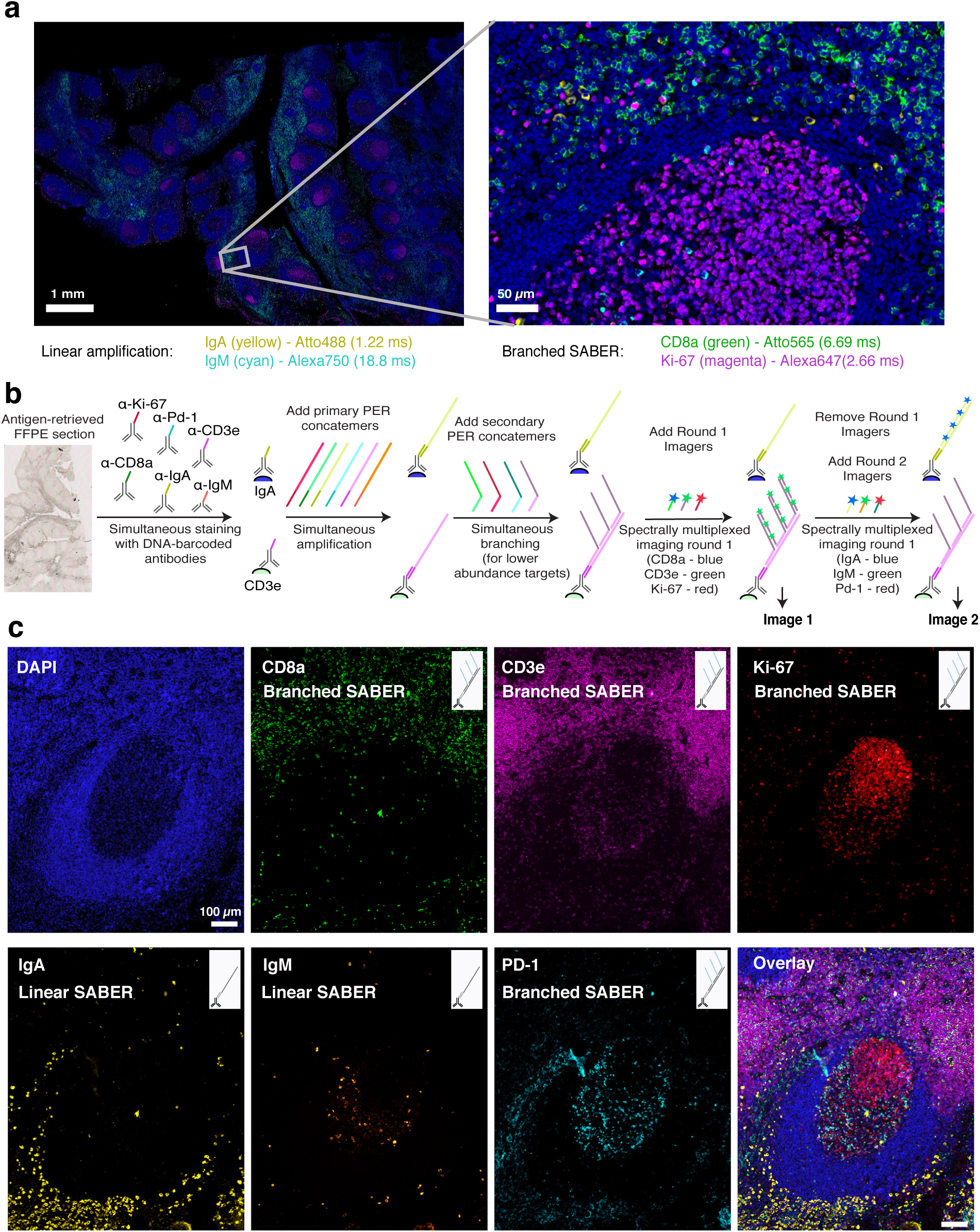
Highly multiplexed imaging in FFPE human tonsil samples. **(a)** Centimeter-scale whole-slide imaging of human tonsil sections with 5-color spectral multiplexing (DAPI + 4 targets). A zoom-in view of the region marked with the grey box showing 4-target imaging with subcellular resolution with a 20× objective and exposure times for each target (auto-exposure setting) is given at the bottom. For IgA and IgM (higher copy number), linear amplification was enough to get high enough signal to achieve auto-exposure times of 1-20 ms under optimized conditions. For Ki-67 and CD8a (lower copy number)^74, 75^ branched amplification (one round of branching) was applied to allow auto-exposure times of 2-10 ms. **(b)** Higher Multiplexing: The schematic for multiplexed imaging workflow where all antibodies are applied simultaneously, followed by simultaneous amplification, and sequential rounds of imaging. IgA and CD3e labeling structures are shown as examples to illustrate the workflow for linear and branched SABER on the same sample. **c)** Images show a zoom-in view of a germinal center in human FFPE tonsil sections imaged in 7-colors (DAPI + 6-targets) with a single exchange round (round 1: top row; round 2: bottom). 4 of the targets CD8a (Atto488), CD3e (Alexa647), Ki-67 (Alexa750) and PD-1 (Alexa750) were visualized with simultaneous branched SABER amplification, whereas IgA (Atto488) and IgM (Atto565) were visualized with linear amplification.

Next, we utilized DNA-Exchange to increase the multiplexing level and imaged different cell types in germinal centers with 6-markers (CD8a, CD3e, Ki-67, PD-1, IgA and IgM) **(Fig. 5b)**. SABER amplification was applied for all targets simultaneously by hybridization of six orthogonal PER concatemers onto antibodies barcoded with the bridge strands. Simultaneous branching was applied to the 4 less abundant targets (CD3e, CD8a, Ki67 and PD-1) following the primary concatemer hybridization, and 3 markers per round were imaged for 2 rounds as depicted in **Fig. 5b**. A germinal center with interfollicular zone is displayed in **Fig. 5c**. With our exchange protocol, it was possible to perform efficient imager removal in 10 min without displacing the concatemers (**Fig. S5)**.

### Highly multiplexed Immuno-SABER imaging in mouse retina cryosections

Next, we validated highly multiplexed Immuno-SABER in thick (40 µm) cryosections of mouse retina and demonstrated ten-target in situ protein imaging. We first screened antibodies against a list of targets that have defined staining patterns, including cone arrestin, SV2, VLP1 (Visinin-like protein 1), Rhodopsin, Calretinin, PKα (Protein kinase C alpha), GFAP (Glial fibrillary acidic protein), Vimentin, Collagen IV, Calbindin. VLP1, Calretinin and Calbindin are all calcium-binding proteins^57, 58^. Calretinin labels a subset of amacrine and ganglion cells. Although it has been suggested that calbindin also exists in amacrine cells and ganglion cells, the antibody targeting calbindin we used mostly labels horizontal cells^58^. Rhodopsin is located in the rod photoreceptors. GFAP labels astrocytes, and Vimentin labels Muller cells^9^. Collagen IV marks blood vessels. PKCα labels blue cone cells and rod bipolar cells^58^. We then conjugated DNA bridge strands to those antibodies, and validated the specificity and affinity of DNA-conjugated antibodies by comparing the staining patterns from conventional staining using unmodified primary antibodies followed by fluorophore secondary antibodies **(Fig. S6a)**.

As a control experiment for DNA-Exchange, we compared pre- and post-washing images for SV2, and found the imager strands were efficiently removed after washing (**Fig. S6b**). To test whether the washing step causes sample damage or signal loss, we imaged SV2 for three rounds and found the correlation coefficient between images was above 0.95, suggesting the washing condition is sufficiently mild and non-disruptive (**Fig. S6c**).

The 40 µm mouse retina section was first incubated with all DNA-conjugated antibodies simultaneously. All SABER concatemers were then added simultaneously to the sample and left at room temperature for overnight, followed by washing and microscopic imaging with sequential addition of fluorophore-attached imager strands. A z-stack of images was acquired for each target, and DAPI was imaged in every exchange cycle to monitor sample drift. The maximum projected images of each stack were computationally aligned using a subpixel registration algorithm using DAPI as the drift marker^9^, and pseudo-colored for presentation **(Fig. 6a)**.

**Figure 6.**
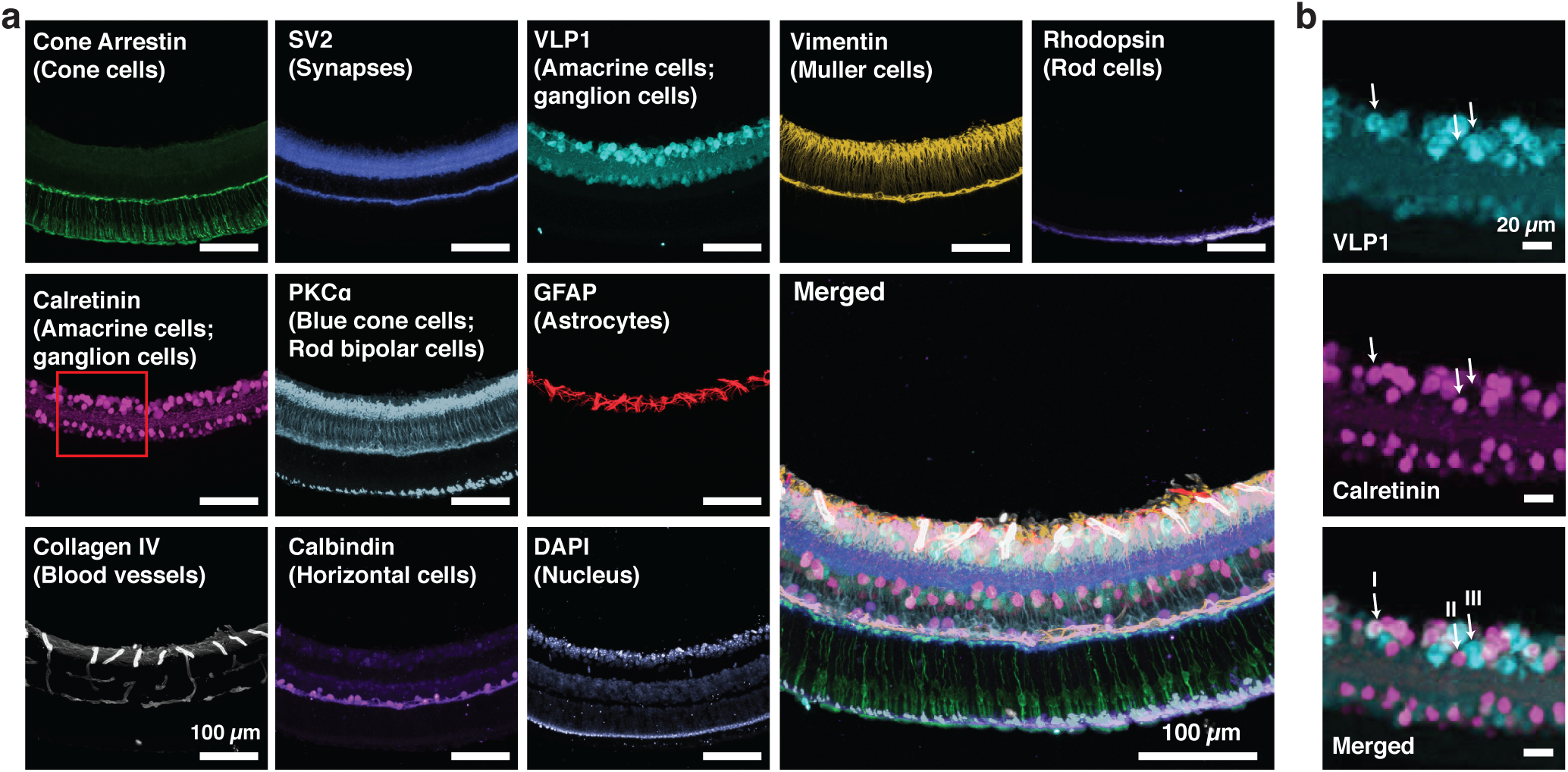
Highly multiplexed protein imaging in mouse retina cryosections. **(a)** Ten protein targets labeling various retinal cell types as noted in the figure were visualized using immuno-SABER in a 40 µm mouse retina cryosections. The images were aligned using the DAPI channel computationally and pseudo-colorized for demonstration. **(b)** Zoom-in view of the area marked by the white rectangle in **a**. Three cell subtypes (marked with arrows, I: VLP1^+^ and Calretinin^+^, II: VLP1^-^ and Calretinin^+^, III: VLP1^+^ and Calretinin^−^) can be differentiated based on VLP1 and Calretinin expression.

The highly multiplexed imaging result showed all targets were successfully captured using Immuno-SABER with expected staining patterns. Unexpectedly, we found that two calcium-binding proteins, VLP1 and Calretinin, together identify three populations of cells, VLP1^+^/Calretinin^-^, VLP1^-^/Calretinin^+^ and VLP1^+^/Calretinin^+^ **(Fig. 6b)**. The antibody specificity was validated by comparing the staining patterns of three different antibodies against the same target **(Fig. S6d-e)**. We also confirmed this result using conventional immunostaining methods with unconjugated primary antibodies followed by fluorophore labeled secondary antibodies **(Fig. S6f)**.

### Rapid, highly multiplexed super-resolution imaging combining immuno-SABER and Expansion Microscopy

The spatial resolution of conventional fluorescence microscopy is limited due to diffraction of light. A variety of super-resolution imaging methods, including Structure Illumination Microscopy (SIM), Stimulated Emission Depletion (STED), localization-based methods (e.g. STORM and PAINT) have been developed to overcome this limitation^59^. Recently, ExM, which improves the practical resolution by physically expanding hydrogel embedded samples, has been gaining popularity especially for imaging of tissues without specialized super-resolution instruments^47^. Up to 20-fold expansion has been achieved, enabling ∼25 nm spatial resolution imaging^60^. However, a key challenge for ExM is that the physical expansion also dilutes fluorescence signals, creating a high need for signal amplification. Both secondary antibodies and HCR have been used to achieve higher signals in ExM with limited multiplexing capability^61^. A technique termed Magnified Analysis of the Proteome (MAP) was developed to achieve higher multiplexing by combining ExM with repeated antibody staining and retrieval^62^. Up to 7-rounds of sequential protein labeling was achieved with this method; however, due to the slow permeation of the antibodies into the thick expanded tissue samples, each round of primary and secondary antibody staining takes 2-9 days, making the approach slow and laborious for high-multiplexing.

We tested Immuno-SABER’s applicability for ExM samples for rapid, highly multiplexed super-resolution imaging. For this purpose, we modified the 5′end of the SABER concatemers with an acrydite moiety that could be incorporated into polyacrylate hydrogels. We first tested the protocol by staining mouse retina cryosections with DNA-conjugated SV2 antibodies **(Fig. 7a)**. The result indicated ∼ 3-fold expansion, which was consistent with the expansion factor achieved in the published exFISH protocol^63^. It was smaller than the expansion factor reported in the original expansion protocol (∼4.5-fold) due to the shrinkage of the gel in the ionic gel re-embedding solution or in the imaging buffer with 0.5× PBS. To further evaluate Immuno-SABER for super-resolution imaging, we imaged a pre-synaptic marker, Bassoon, and a post-synaptic marker, Homer1b/c, in fixed primary mouse hippocampal neuron culture with and without expansion **(Fig. 7b** and **Fig. S7a)**. While Bassoon and Homer1b/c were readily separated after expansion, they strongly overlapped without expansion. We next demonstrated multiplexed super-resolution imaging by visualizing six targets (Vimentin, Collagen IV, Rhodopsin, Calretinin, GFAP, SV2) in mouse retina cryosections **(Fig. 7c)**. Since the expanded sample was reaching ∼350 µm in thickness we increased both the incubation time and wash time for imagers (to 45 min and 1.5 h respectively) to achieve optimal imaging quality. Under these conditions, we were able to perform six-target imaging using two rounds of imager incubation in six hours, which is substantially faster than the MAP protocol that would take >3 days^62^. To further increase the speed, we incorporated an alternative fluorophore removal protocol that shortened the removal time from 1.5 h to only 10 min. In this alternative protocol, a disulfide bond was included in the imager strands between the fluorophore and DNA sequences. Hence, the fluorescence signal could be quickly displaced using reducing agents (e.g. TCEP)^64^ (**Fig. S7b** and **c**).

**Figure 7.**
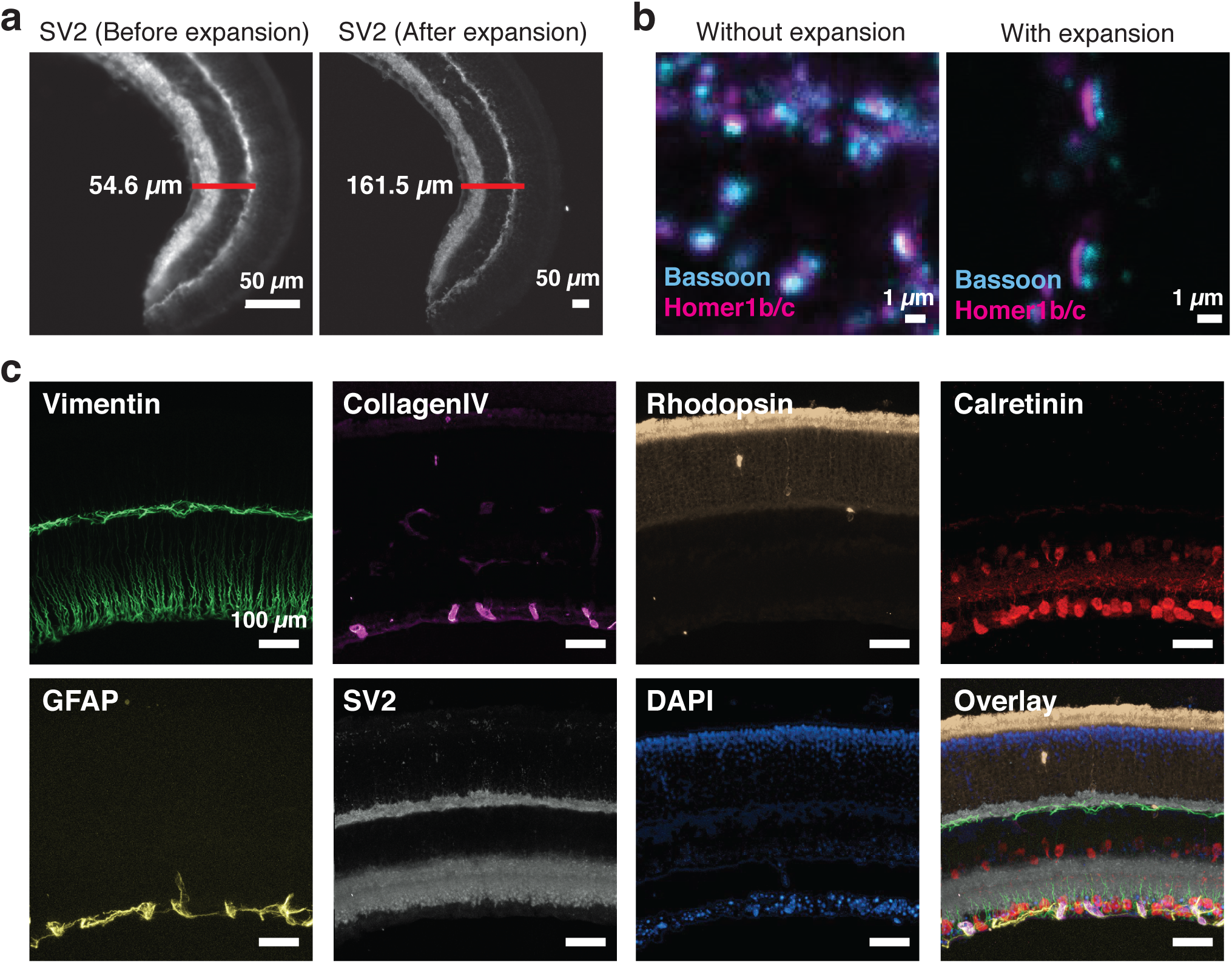
Highly multiplexed super-resolution imaging using Immuno-SABER-Expansion Microscopy. **(a)** Before and after expansion imaging of SV2 in mouse retina sections. **(b)** Images of pre- and post-synaptic sites of neuronal synapses in a fixed primary mouse hippocampal neuron culture with and without expansion (different fields of view are shown). **(c)** Super-resolution imaging of six protein targets in the originally 40 µm-thick mouse retina section (expanded ∼3-folds). Two rounds of DNA-Exchange were performed to visualize all six targets. DAPI was imaged in both rounds to serve as a registration marker. The images were maximum projections of z-stacks. They were drift-corrected using DAPI channels, and pseudo-colored for presentation.

## Discussion

Mapping molecular heterogeneity at single cell level is of great interest, and a number of large-scale collaborative projects, such as the Human Cell Atlas Project and the Human BioMolecular Atlas Program, have been launched to develop technologies for such efforts^65, 66^. In the past few years, advancement of single cell DNA/RNA sequencing of dissociated cells has enabled researchers to discover new cell types and analyze cell population change. However, the loss of spatial information makes it difficult to understand how different cell types interact with each other to realize biological functions. Hence, *in situ* analysis with highly multiplexed microscopy imaging methods such as Fluorescence in situ Sequencing (FISSEQ)^15^, Multiplexed Error-robust FISH (MERFISH)^64, 67^, Sequential Barcoding FISH (seqFISH)^18, 41^, Spatially-resolved Transcript Amplicon Readout Mapping (STARmap)^17^ and ClampFISH^68^ are poised to recover the molecular identities of cells in their spatial context by allowing high levels of multiplexing. As we recently demonstrated, SABER amplification is also compatible with FISH-based multiplexed *in situ* visualization of RNA and DNA molecules with signal amplification^46^.

Here, we focus on highly multiplexed *in situ* protein imaging, an application that is relatively less developed and in high need of new innovations. We previously reported the DNA-Exchange-Imaging (DEI) technique, where we first demonstrated multiplexing through barcoding of antibodies with orthogonal DNA sequences for simultaneous staining followed by *in situ* DNA detection by hybridization^9, 12^. To achieve reliable signal to noise, DEI largely relied on secondary antibodies from non-overlapping species, which is a hard to satisfy requirement for scaling up. With Immuno-SABER we provide a new multiplexed amplification capability to increase the detection sensitivity and tune it independently for multiple targets, which is also instrumental to increase the imaging speed by decreasing the need for long exposure/scanning times. Amplification is crucial for high sensitivity to enable confident detection of rare targets that otherwise can’t sufficiently accumulate enough fluorophores and yield low signal-to-noise especially in the tissue samples. Utilizing Immuno-SABER with primary antibodies, we demonstrate as much as 5-20-fold signal enhancement for linear SABER and ∼50-fold for branched SABER (>5-fold higher signal than fluorophore-conjugated secondary antibodies), and ∼180-fold for iterative SABER using primary antibodies in tissues. Through the use of orthogonal concatemers, we validate simultaneous amplification for up to 10 protein targets in the same sample. For designing these sequences, we rely on a 3-letter code (where G nucleotides are avoided in the primer sequence) and an efficient and non-leaky stopper for the polymerase to achieve controlled signal amplification through PER^45, 46^. We performed *in situ* testing of a pool of 32 from our 50 *in silico* designed sequences^46^, where we detected minor crosstalk for only 1 of the 1024 pairs (32×32) under our experimental conditions. An even larger pool of PER sequences have been *in silico* designed and are readily available^46^. Owing to the simplicity of sequence design and minimal sequence constraints PER requires, it is straightforward to scale up the orthogonal sequence pool size and reach much higher multiplexing levels for detailed protein maps.

From a practical perspective, Immuno-SABER allows improvement of throughput in two ways. Firstly, in contrast to methods involving repeated antibody staining, Immuno-SABER allows application of multiple primary antibodies in one step. Also the orthogonal amplification concatemers are applied at once, and multiplexed imaging is achieved by rapid exchange of short imager strands. Antibody incubation is usually the longest part of the immunostaining protocol, especially for thicker tissue sections. When this step is performed at every cycle of sequential labeling, sample preparation time increases substantially. Secondly, signal amplification by Immuno-SABER allows substantial shortening of image acquisition times. Methods such as DEI and CODEX that operate without signal amplification may necessitate long (hundreds of ms) camera exposure times per frame to collect enough photons. This is particularly problematic for large-scale experiments such as tissue mapping, high content screening or whole slide scanning where centimeter scale areas need to be imaged and imaging time is the primary bottleneck. With Immuno-SABER, it is possible to image such large samples at a significantly faster rate enabled by shorter exposure times (we were able to operate at <10 ms exposure times for markers like CD8a, Ki-67 and IgA in human tonsil FFPE samples using a commercial slide scanner at subcellular resolution with a 20× objective).

As demonstrated here the ability to preprogram and control the amplification fold for each target (in this case through application of linear or branched SABER on the same section) is very valuable for multiplexed imaging of complex tissue samples with target proteins expressed at vastly variable levels. The 10-100 fold improvement in image acquisition speed^46^ can deliver 1-2 orders of magnitude enhancement of throughput, which is much needed for discovery-oriented cell/tissue profiling and large-scale mapping projects. The resulting high-dimensional Immuno-SABER data can then be processed and analyzed with existing tools such as Histocat^69^, Wanderlust^70^, t-SNE^71^, or viSNE^72^, utilized for mass cytometry or cyclic immunostaing methods like t-CycIF^8^.

In addition to the abovementioned features, Immuno-SABER is also compatible with widely available microscopy platforms, including standard wide-field and confocal microscopes or slide scanners. Since the exchange part can be performed off the imaging platform (while another sample is being imaged), integration of a fluidic system is optional for higher automation. Sample preparation can also be automated since the protocols compatible with standard workflows would allow the incorporation of existing automated stainers. This compatibility promises easy adoption by a broad range of researchers (from laboratories performing exploratory or small-scale preparations without dedicated instruments to specialized laboratories with high automation looking to improve the throughput).

Furthermore, Immuno-SABER can be of great interest to the super-resolution community in need of multiplexed tissue imaging experiments. Through combination with expansion microscopy, we demonstrate here a highly multiplexed super-resolution application where we imaged 6 targets in tissue samples expanded ∼3-fold in three dimensions within six hours in contrast to ∼4-5 days using previously published protocols such as MAP^62^. With the alternative protocol we provide, the experimental time can be further reduced to 3 hours. Here, besides the substantial improvement in speed, signal amplification is critically needed as expansion naturally decreases the density of the target molecule. This hampering decline of fluorescence signal can be prevented through Immuno-SABER.

In summary, we offer a simple and effective method for highly multiplexed and sensitive *in situ* protein detection that enables individually programmable signal amplification for each target. With these capabilities and the demonstrated applicability for diverse samples, Immuno-SABER is poised to provide higher-throughput for imaging assays ranging from nanometer-scale super-resolution studies to centimeter scale tissue mapping efforts. We expect Immuno-SABER would be useful to a broad range of researchers and would prove valuable for a wide spectrum of potential applications including tissue atlases, tumor/disease profiling, *in situ* single-cell validation for bulk assays such as flow or mass cytometry, or CITE-Seq^73^, and complementing single-cell RNAseq analysis with the functional information of protein expression. Being principally compatible with standard workflows and equipment used for clinical specimen, Immuno-SABER can also be utilized for pathology and, biomarker screening and discovery. Along with other single-cell and *in situ* ‘omics’ methods, we believe it will greatly contribute to next generation multi-dimensional analyses of biological samples.

## Acknowledgements

We thank Peter Sorger, Zoltan Maliga, and Constance Cepko for discussion and critical input. We thank the Neurobiology Department and the Neurobiology Imaging Facility for consultation and instrument availability that supported this work. This facility is supported in part by the Neural Imaging Center as part of an NINDS P30 Core Center grant #NS072030. We thank Maël Manesse, Tonora Archivald and David Bowman for help with the FFPE samples. We thank Shanshan Wang for providing neuronal cultures.

## Data and Software Availability

The data and essential custom scripts for image processing and plotting will be made available from the corresponding authors P.Y., S.K.S., and Y.W. upon request.

## Supplementary Information

Supplementary Information contains seven supplementary figures, detailed experimental materials and methods, and supplementary references.

