## Supplemental Information for "Highly multiplexed *in situ* protein imaging with signal amplification by Immuno-SABER"

[1] Wyss Institute for Biologically Inspired Engineering, Harvard University, Boston, MA 02115, USA
[2] Department of Systems Biology, Harvard Medical School, Boston, MA 02115, USA
[3] Department of Genetics, Harvard Medical School, Boston, MA 02115, USA
[4] Department of Biomedical Informatics, Harvard Medical School, Boston, MA 02115, USA
[5] Image and Data Analysis Core, Harvard Medical School, Boston, MA 02115, USA
[6] Laboratory of Systems Pharmacology, Harvard Medical School, Boston, MA 02115, USA
[7] Media Lab, Massachusetts Institute of Technology (MIT), Cambridge, MA 02139, USA
[8] Department of Neurobiology, Harvard Medical School, Boston, MA 02115, USA
[9] Pathology Department, Beth Israel Deaconess Medical Center, Boston, MA 02115, USA
[10] Present address: Department of Genome Sciences, University of Washington, Seattle, WA 98195, USA
[11] These authors contributed equally.
[12] Lead contact

### Contents

**List of Supplementary Figures**
Supplementary Figure S1 (related to Fig. 1)
Supplementary Figure S2 (related to Fig. 2)
Supplementary Figure S3 (related to Fig. 3)
Supplementary Figure S4 (related to Fig. 4)
Supplementary Figure S5 (related to Fig. 5)
Supplementary Figure S6 (related to Fig. 6)
Supplementary Figure S7 (related to Fig. 7)

**Materials and Methods**
1. Preparation of SABER extensions
2. Antibody conjugation and purification
3. Microtubule staining in cell culture, imaging FWHM estimation
4. Preparation, staining, SABER application, quantification and analysis on human tonsil FFPE sections
5. Preparation, staining, SABER application and quantification on mouse retina cryosections
6. Preparation, staining, SABER application on whole mount retina samples
7. Crosstalk analysis in cell culture
8. Expansion microscopy

**List of Tables**
Table S1. Optimized concatemer extension conditions and sequences for the primer library
Table S2. Imager strands used for visualization
Table S3. Bridge sequences
Table S4. Antibodies used in SABER experiments and conjugated sequences

**Supplementary References**

### Supplementary Figures

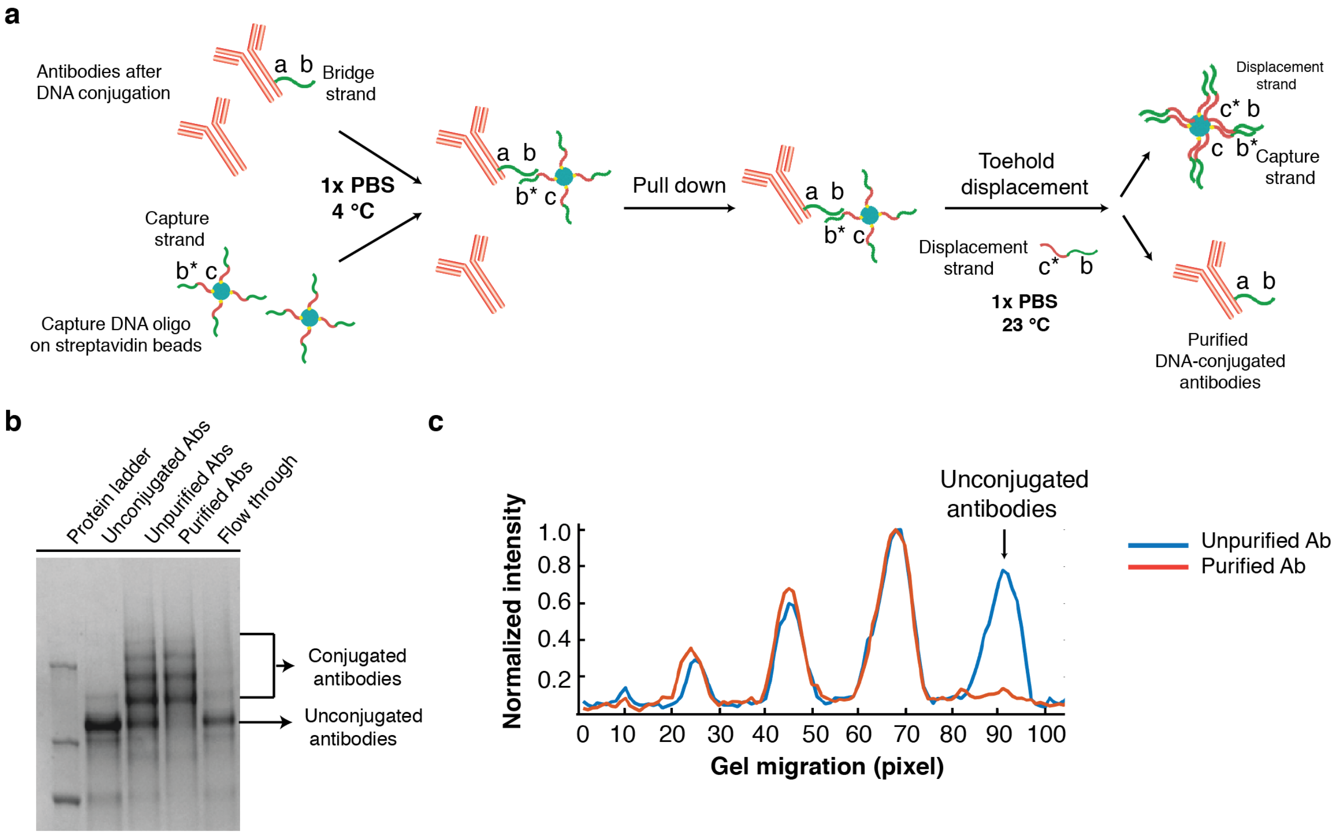


**Supplementary Figure S1 (related to Fig. 1). Optional purification of DNA-conjugated antibodies using a toehold displacement-mediated DNA affinity pull-down. (a)** Schematic of the pull down assay. Biotin-modified capture DNA strands (biotin-*c*-*b**, where *c* is 15-nucleotides and *b** is 16-nucleotides and) are attached to streptavidin beads, and used to pull down DNA-conjugated antibodies after antibody-bridge DNA conjugation (bridge strand sequence: *a-b,* where *b* is 16 nucleotide*s*). The attached antibodies are dissociated from the beads using toehold displacement strands (*c*-b*) that compete with capture strands[1] on antibodies. While optional for Immuno-SABER, we found that purifying the DNA-conjugated antibodies via pull-down and toehold-mediated displacement may be helpful to improve the signal for select antibodies. **(b)** Visualization of purification products using an SDS-PAGE gel assay. After DNA conjugation, the majority of antibodies were conjugated with 0, 1 or 2 DNA oligos per antibody. After purification, antibodies without DNA were removed. **(c)** Plot of protein densities for the bands in **b**. The band corresponding to removed unconjugated antibodies is marked with an arrow.

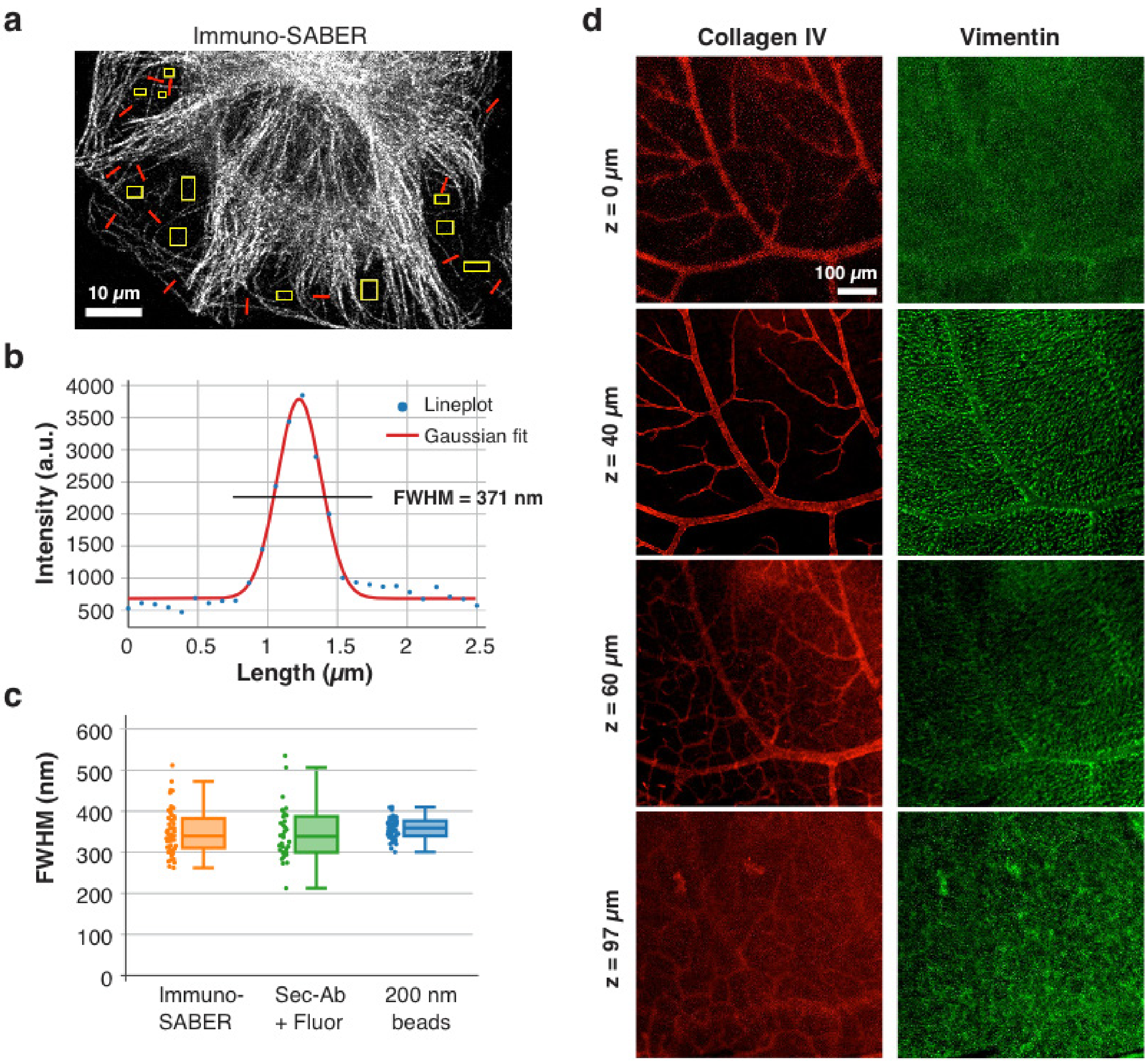


**Supplementary Figure S2 (related to Fig. 2). Resolution and penetration controls for Immuno-SABER. (a)** ~2.5 μm long lineplots (red) were made over the thin tubules to estimate the observed resolution. Yellow-boxed background regions were used to estimate the background for subtraction. **(b)** A typical line plot along a tubule and the Gaussian fit, where full-width-half-maximum (FWHM) was calculated as 371 nm. **(c)** Mean FWHM values were calculated for 30-45 lineplots from cells stained with Immuno-SABER or fluorophore-conjugated secondary antibodies samples and the distribution was displayed as a box plot. A similar calculation was performed for 200 nm fluorescent beads. Mean FWHM was not significantly different (p value is 0.360 for Immuno-SABER and 0.335 for conventional secondary antibody staining). **(d)** Visualization of Collagen IV and Vimentin at multiple depths of the whole mount mouse retina shown in **Fig. 2f**. Selected confocal planes are shown. Vimentin stains the Muller cells and Collagen IV stains the blood vessels, both of which are localized predominantly in the segments from nerve fiber layer to outer plexiform layer (~100 μm)[2].

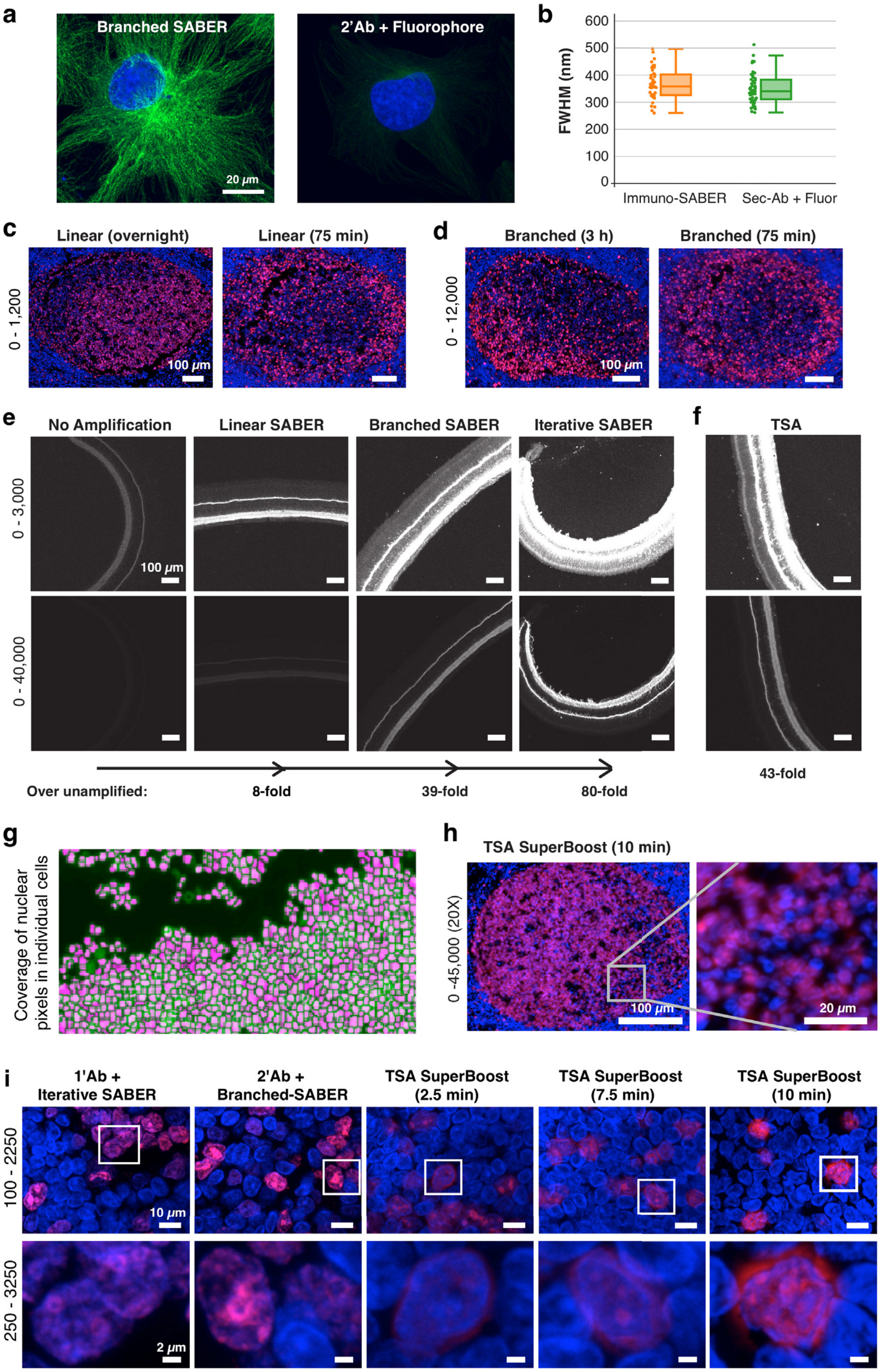


**Supplementary Figure S3 (related to Fig. 3). Exponential amplification via branching. (a)** Alpha-tubulin staining (Alexa647) in cultured BS-C-1 cells (max projections from confocal z-stack). **(b)** Mean FWHM values were calculated for 43 lineplots from cells stained with branched Immuno-SABER. For comparison values for conventional staining is also included. Mean FWHM was not significantly different than the sub-diffraction 200 nm bead samples (plotted in **Fig. S2c**). **(c-d)** Images display a typical germinal center in human FFPE tonsil samples stained for Ki-67 (Alexa647, red) by Immuno-SABER. DAPI stain (blue) is shown for reference. Similar amplification levels were obtained by long and short hybridization times (at 37°C) for the primary concatemer and branching concatemer (75 min each). **(e)** Iterative SABER of SV2 (Alexa647) in a 40 µm mouse retina cryosection. **(f)** For comparison SV2 staining with TSA was performed using mono HRP conjugated secondary antibodies. **(g)** Watershed-based segmentation was used to segment (pink) the pixels corresponding to nuclei of each individual cell after deep-learning identification of nuclear contours, as in **Fig. 3f. (h)** Application of tyramide-Alexa647 for the maximum recommended incubation of 10 min. The germinal center image is scaled in the same range with the top row of **Fig. 3j**. Zoom-in on the right is included to display the significant blurring of the signal at 10 minutes incubation. **(i)** Zoom-in views of the in **Fig. 3j** displayed at different scaling ranges for comparative visualization.

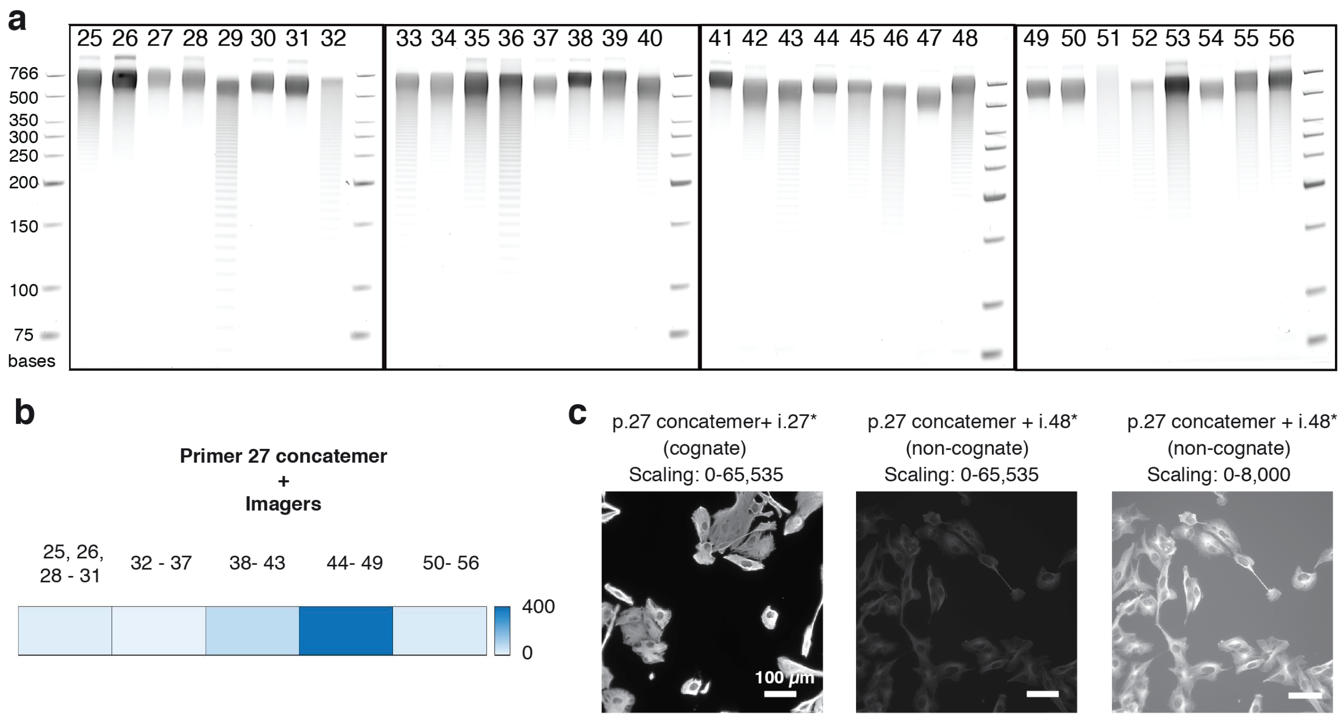


**Supplementary Figure S4 (related to Fig. 4). Primer characterization and crosstalk analysis. (a)** Full PAGE gels used to crop the bands displayed in **Fig. 4a**. Qualitatively, 18 of the 32 sequences (such as #29) displayed a broader distribution with a ladder of shorter products visible albeit these bands being much dimmer than the predominant concatemer band. Although being extended, one sequence (#51) had a more even distribution in the upper length regime without a clear predominant band. Based on these distributions, particular applications may favor a subset of the sequence library, i.e. high efficiency primers (for example primers 27, 28, 30, 31, 37, 38, 41, 44, 47, 49, 50, 54) may be more favorably utilized for more quantitative experiments or for lower abundance proteins. **(b)** Crosstalk analysis of primer sequence p.27. BS-C-1 cells were fixed and stained with DNA-conjugated antibodies targeting alpha-Tubulin. Concatemers extended from primer sequence p.27 were hybridized to the antibodies, followed by addition of individual imager strands. Only the imager strand Alexa647-i.48* showed non-negligible crosstalk.

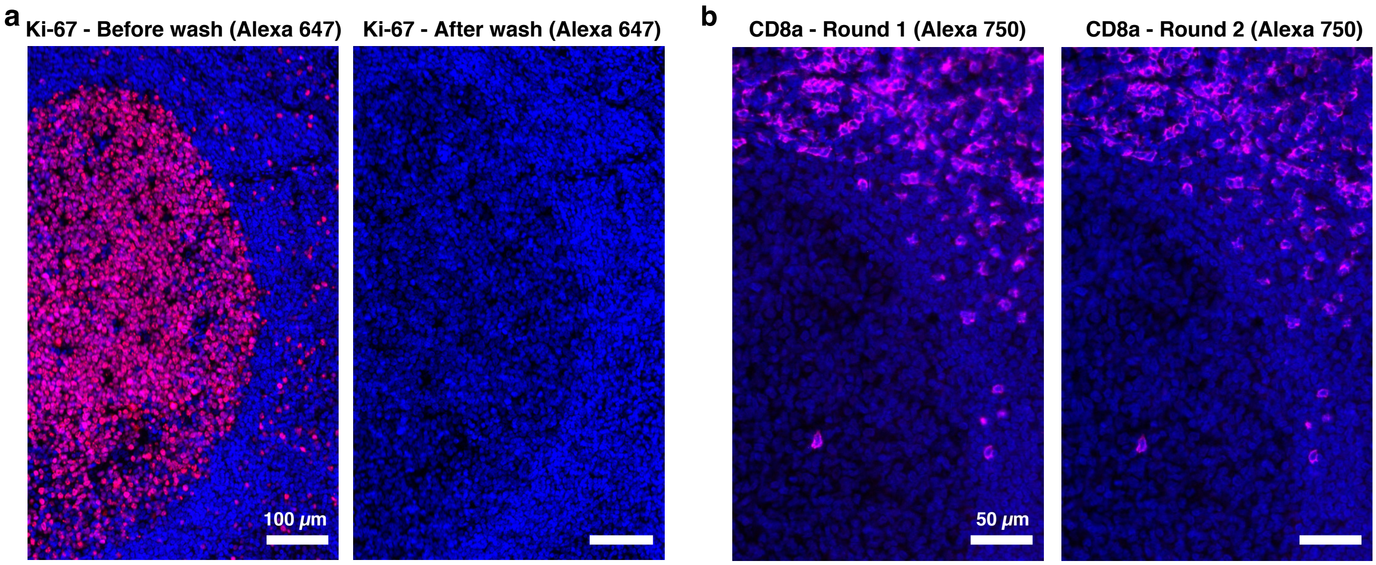


**Supplementary Figure S5 (related to Fig. 5). Controls for exchange imaging of FFPE human tonsil sections.**
**(a)** Imagers can efficiently removed with a 10 min wash using 50% formamide in PBS, as shown in before and after wash images of the same section stained for Ki-67 (red) with linear amplification, imaged and displayed under the same conditions. DAPI stain is shown in blue. **(b)** Under this wash condition (10 min with 50% formamide in PBS at room temperature) the concatemers are not displaced, as shown by exchange imaging of CD8a by re-binding the imagers to the same target and re-imaging the tissue section (linear amplification).

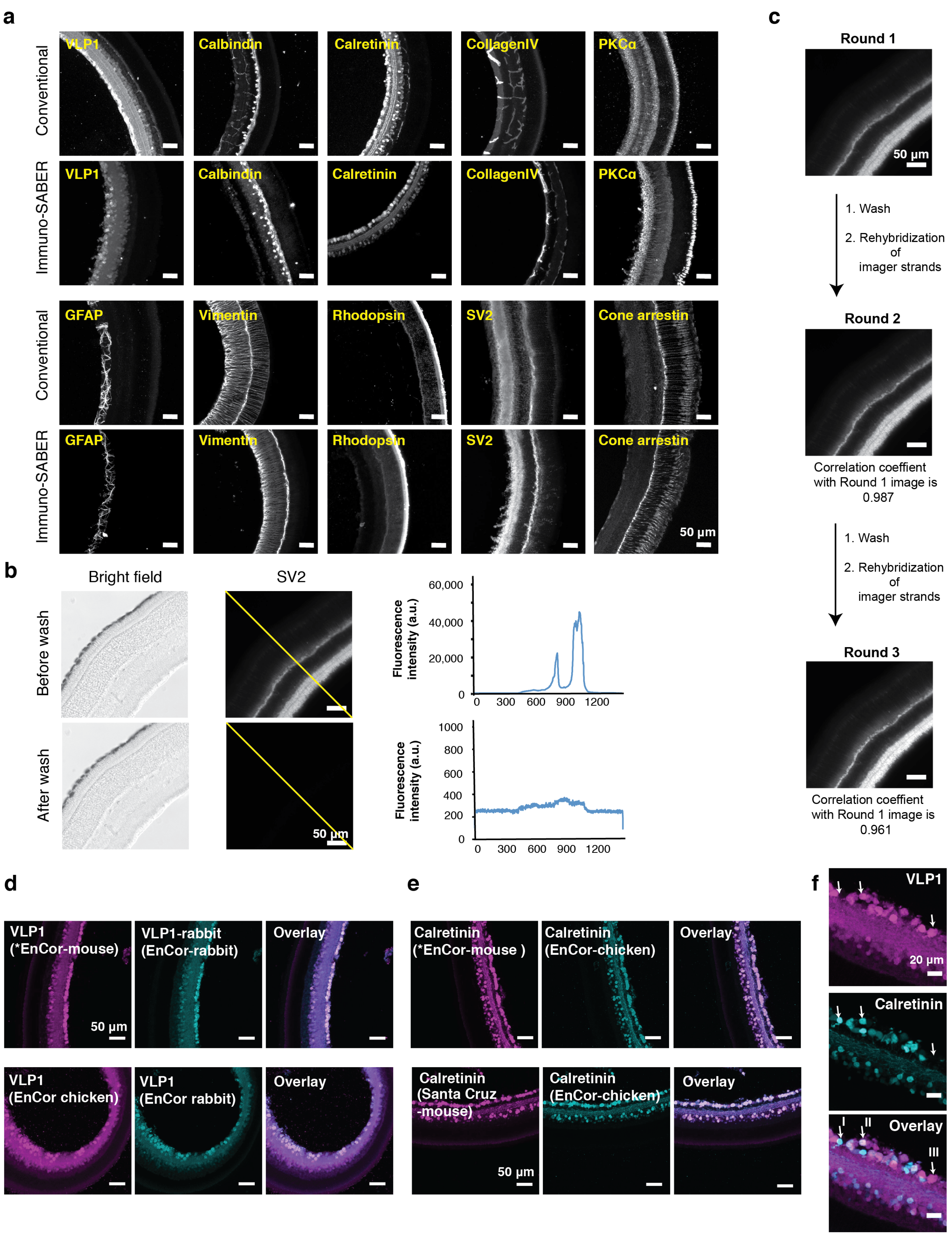


**Supplementary Figure S6 (related to Fig. 6). Control experiments for highly multiplexed imaging in mouse retina sections. (a)** Comparison of antibody staining patterns before and after DNA conjugation. The images for unconjugated antibodies were taken using conventional fluorophore-conjugated secondary antibody labeling, and the images for antibodies after conjugation were taken using primary antibody-Immuno SABER labeling (images are displayed at individual contrast levels). **(b)** Efficiency of washing to remove the imager strands. A 30 µm mouse retina section was stained with DNA-conjugated SV2 antibodies and imaged using Alexa647-i.26* imager with a widefield microscope. Imager strands were washed using 0.1× PBS with 30% formamide at room temperature for 3×10 minutes. Before and after images were taken using the same imaging setting. The fluorescence intensity of indicated yellow line was measured using FIJI[3]. **(c)** Washing conditions maintained sample integrity without signal loss. SV2 in mouse retina sections was imaged for three rounds and the correlation coefficient was calculated between the images. **(d-e)** Validation of the VLP1 and calretinin antibodies with conventional indirect immunostaining: The VLP1 (EnCor-mouse) and calretinin (EnCor-mouse) that were used for the multiplexing experiments (marked with asterisk) in **Fig. 6** were tested for specificity by co-staining with other antibodies targeting the same targets, followed by visualization using fluorophore-conjugated secondary antibodies. **(f)** Three cell subtypes (marked with arrows, I: VLP1$^+$ and Calretinin$^+$, II: VLP1$^-$ and Calretinin$^+$, III: VLP1$^+$ and Calretinin$^-$) identified in the multiplexed mouse retina imaging experiment were verified using conventional immunostaining with unconjugated primary antibodies (VLP1 by EnCor-rabbit and calretinin by EnCor-mouse) and fluorophore secondary antibodies.

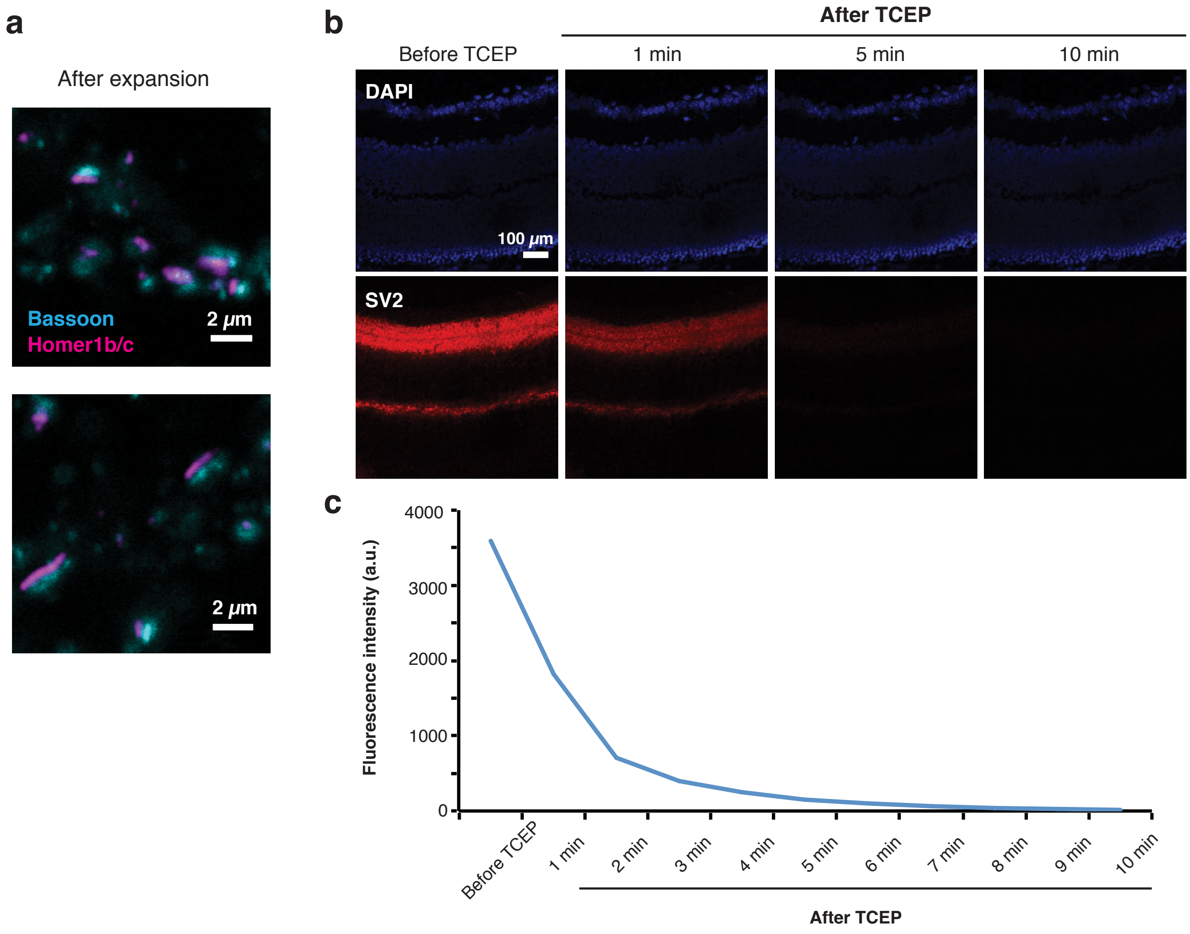


**Supplementary Figure S7 (related to Fig. 7). Other sample orientations and alternative reduction-mediated fluorophore removal timeline. (a)** Additional post-expansion images of neuronal synapses showing different orientations. **(b)** Alternative fluorophore removal strategy using TCEP reduction in thick expanded samples using TCEP. A mouse retina section stained with SV2 was expanded and visualized with disulfide bond modified imager strands. The fluorophores on the imager strands were cleaved using TCEP reduction. The fluorescence signal was monitored in a time course for 10 min. **(c)** Quantification of fluorescence signals in **b** before and after TCEP reduction.

### Materials and Methods

#### 1. PER sequences and preparation of SABER concatemers

***In vitro* extension of primers.** Concatemer extensions were prepared as described previously[4]. Typically, 100 µl reactions were prepared in PBS with final concentrations of: 10 mM $MgSO_4$, 400-1000 units/ml of Bst LF polymerase (NEB M0275L or McLab BPL-300), 600 µM each of dATP/dCTP/dTTP (NEB #M0275L), 100 nM of Clean.G (5'-CCCCGAAAGTGGCCTCGGGCCTTTTGGCCCGAGGCCACTTTCG-3') hairpin[5], 50 nM-1.5 µM hairpin, and water to 90 µl. The Clean.G hairpin has a 5' stretch of C's. Pre-incubation with Clean.G helps to get rid of the impurities in the dHTP mixtures (made up of dATP, dTTP, and dCTP), which may have small amounts of dGTP contamination[5]. After incubation for 15 min at 37°C, 10 µl of 10 µM primer was added, and the reaction was incubated another 1-3 h at 37°C followed by 20 min at 80°C to heat inactivate the polymerase. Reaction products can be stored at -20°C for several months. In our demonstrations, PER products were diluted into concatemer hybridization solutions for binding to the bridge sequences. Alternatively, concatemers can be purified and concentrated using a MinElute (Qiagen #28004) kit with distilled water elution to reduce volume and salt concentration from the reaction condition. Primer sequences and details of the extension conditions utilized for **Fig. 4** are listed in **Table S1**. These primer sequences were presented as the 3' tail of the complements of the bridge strands given in **Table S3,** in the format: 5'- bridge*- tt (spacer nucleotides) - 9mer primer sequence - 3', where * denotes the reverse complement. For the extensions in **Fig. 4**, the 25mer-tester* bridge was used for all the primers. The strands were obtained from IDT. Primers were synthesized and provided with standard desalting. Hairpins were ordered with 3' inverted dT modification to ensure they cannot be extended. Due to the modification, they were ordered with HPLC purification. Details of the primer and hairpin design criteria are described in our previous work[4, 5]. For primary concatemers, we utilize concatemers reaching the 600-700 base based on empirical experience[4].

***Gel electrophoresis.*** After extension, for internal quality control lengths of concatemers were evaluated by diluting 1 µl of *in vitro* reaction with 19 µl water. For quality controlling, samples were then run on 1-2% E-Gel EX agarose gels (Thermo Fisher #G402001) for 10 min on the E-gel apparatus (Invitrogen, iBase) alongside a 1 kb Plus DNA Ladder (Invitrogen) and imaged with the Sybr Gold channel on a Typhoon FLA 9000 scanner.

For the comparison gel in **Fig. 4** and **Fig. S4**, unpurified concatemers were run using 6% TBE-UREA PAGE gels (Thermo Fisher) at 55°C. The gel was pre-run for 1 h before loading the samples. 160 ng Quick-Load Purple Low Molecular Weight DNA Ladder (NEB #N0557S) was loaded as size reference. The reaction products were diluted 1:7 with 2× Urea-Loading Dye, and denatured at 95°C for 5 min. 9 µl from each sample was loaded on the gel. Both samples and the ladder were denatured. Samples were ran for 20 min at 75 V, and at 130 V for 1 h. Gels were stained with 1:10,000 SybrGold in 0.5× Tris-Borate-EDTA (TBE) for 30 min and scanned on a Typhoon FLA 9000 scanner.

**Table S1. Optimized concatemer extension conditions and sequences for the primer library (for Fig. 4).**

| Primer ID | Primer sequence | Hairpin ID | Hairpin sequence | Hairpin concentration | Time |
|---|---|---|---|---|---|
| p.25 | CCAATAATA | h.25.25 | ACCAATAATAGGGCCTTTTGGCCCTATTATT GGTTATTATTGG/3InvdT/ | 1.5 µM | 3 h |
| p.26 | ATAAACCTA | h.26.26 | AATAAACCTAGGGCCTTTTGGCCCTAGGTTT ATTTAGGTTTAT/3InvdT/ | 9 µM | 3 h |
| p.27 | CATCATCAT | h.27.27 | ACATCATCATGGGCCTTTTGGCCCATGATG ATGTATGATGATG/3InvdT/ | 0.75 µM | 3 h |
| p.28 | CAACTTAAC | h.28.28 | ACAACTTAACGGGCCTTTTGGCCCGTTAAG TTGTGTTAAGTTG/3InvdT/ | 3 µM | 2 h |
| p.29 | TCTAAAATC | h.29.29 | ATCTAAAATCGGGCCTTTTGGCCCGATTTTA GATGATTTTAGA/3InvdT/ | 15 µM | 3 h |
| p.30 | AATACTCTC | h.30.30 | AAATACTCTCGGGCCTTTTGGCCCGAGAGT ATTTGAGAGTATT/3InvdT/ | 5 µM | 2 h |

| | | | | | |
|---|---|---|---|---|---|
| p.31 | TTATTCACT | h.31.31 | ATTATTCACTGGGCCTTTTGGCCCAGTGAAT AATAGTGAATAA/3InvdT/ | 8.5 µM | 2 h |
| p.32 | CTTTTTTTC | h.32.32 | ACTTTTTTTCGGGCCTTTTGGCCCGAAAAAA AGTGAAAAAAAG/3InvdT/ | 15 µM | 3 h |
| p.33 | CCTTCTATT | h.33.33 | ACCTTCTATTGGGCCTTTTGGCCCAATAGAA GGTAATAGAAGG/3InvdT/ | 5 µM | 2 h |
| p.34 | CTCTACTAC | h.34.34 | ACTCTACTACGGGCCTTTTGGCCCGTAGTAG AGTGTAGTAGAG/3InvdT/ | 4 µM | 2 h |
| p.35 | TAAAAACTC | h.35.35 | ATAAAAACTCGGGCCTTTTGGCCCGAGTTTT TATGAGTTTTTA/3InvdT/ | 15 µM | 3 h |
| p.36 | AACTAATCT | h.36.36 | AAACTAATCTGGGCCTTTTGGCCCAGATTA GTTTAGATTAGTT/3InvdT/ | 10 µM | 2 h |
| p.37 | TTTCTCTTC | h.37.37 | ATTTCTCTTCGGGCCTTTTGGCCCGAAGAGA AATGAAGAGAAA/3InvdT/ | 8.5 µM | 2 h |
| p.38 | AACATACTA | h.38.38 | AAACATACTAGGGCCTTTTGGCCCTAGTAT GTTTTAGTATGTT/3InvdT/ | 5 µM | 2 h |
| p.39 | TTCATTTAC | h.39.39 | ATTCATTTACGGGCCTTTTGGCCCGTAAATG AATGTAAATGAA/3InvdT/ | 10 µM | 2 h |
| p.40 | ATCCTACAA | h.40.40 | AATCCTACAAGGGCCTTTTGGCCCTTGTAG GATTTTGTAGGAT/3InvdT/ | 9 µM | 2 h |
| p.41 | CAATCAAAA | h.41.41 | ACAATCAAAAGGGCCTTTTGGCCCTTTTGAT TGTTTTTGATTG/3InvdT/ | 4.5 µM | 3 h |
| p.42 | CTTACAAAC | h.42.42 | ACTTACAAACGGGCCTTTTGGCCCGTTTGTA AGTGTTTGTAAG/3InvdT/ | 5 µM | 2 h |
| p.43 | ACAAATAAC | h.43.43 | AACAAATAACGGGCCTTTTGGCCCGTTATTT GTTGTTATTTGT/3InvdT/ | 5 µM | 2 h |
| p.44 | TTTTCTACC | h.44.44 | ATTTTCTACCGGGCCTTTTGGCCCGGTAGAA AATGGTAGAAAA/3InvdT/ | 4.5 µm | 3h |
| p.45 | CCCTTATTT | h.45.45 | ACCCTTATTTGGGCCTTTTGGCCCAAATAAG GGTAAATAAGGG/3InvdT/ | 4 µM | 3 h |
| p.46 | TCTTTCATT | h.46.46 | ATCTTTCATTGGGCCTTTTGGCCCAATGAAA GATAATGAAAGA/3InvdT/ | 4.5 µM | 3 h |
| p.47 | TTCTTACTC | h.47.47 | ATTCTTACTCGGGCCTTTTGGCCCGAGTAAG AATGAGTAAGAA/3InvdT/ | 8.5 µM | 1 h |
| p.48 | CCATAAATC | h.48.48 | ACCATAAATCGGGCCTTTTGGCCCGATTTAT GGTGATTTATGG/3InvdT/ | 4 µM | 3 h |
| p.49 | CATTTATCC | h.49.49 | ACATTTATCCGGGCCTTTTGGCCCGGATAA ATGTGGATAAATG/3InvdT/ | 6.5 µM | 1 h |
| p.50 | ATACTTCAC | h.50.50 | AATACTTCACGGGCCTTTTGGCCCGTGAAG TATTGTGAAGTAT/3InvdT/ | 4 µM | 1 h |
| p.51 | TACCTCTAA | h.51.51 | ATACCTCTAAGGGCCTTTTGGCCCTTAGAG GTATTTAGAGGTA/3InvdT/ | 6 µM | 3 h |
| p.52 | CTCCTATTT | h.52.52 | ACTCCTATTTGGGCCTTTTGGCCCAAATAGG AGTAAATAGGAG/3InvdT/ | 3 µM | 2 h |
| p.53 | CTATCCAAA | h.53.53 | ACTATCCAAAGGGCCTTTTGGCCCTTTGGAT AGTTTTGGATAG/3InvdT/ | 2 µM | 3 h |
| p.54 | ATCCCTATC | h.54.54 | AATCCCTATCGGGCCTTTTGGCCCGATAGG GATTGATAGGGAT/3InvdT/ | 1 µM | 3 h |
| p.55 | TCATTACTT | h.55.55 | ATCATTACTTGGGCCTTTTGGCCCAAGTAAT GATAAGTAATGA/3InvdT/ | 6.5 µM | 3 h |
| p.56 | CTAAATCTC | h.56.56 | ACTAAATCTCGGGCCTTTTGGCCCGAGATTT AGTGAGATTTAG/3InvdT/ | 3.5 µM | 3 h |
| Test-primer[5] | TCTCTTATT | h.test | ATCTCTTATTGGGCCTTTTGGCCCAATAAGA GATAATAAGAGA/3InvdT/ | Not included in the gel assay | |

The full set of 50 designed primer sequences are available in our recent work[4].

***Imagers.*** SABER imagers are 20mer DNA oligonucleotides with fluorophores on the 5' end and inverted dT modification on the 3' end. Imagers were designed to bind the dimers of the primer unit sequence to achieve stable but easily reversible binding that is necessary for DNA-exchange-imaging. Hence, format of the imager sequences are: 5' - Fluorophore- tt - primer* - t - primer* - t - Inverted dT - 3' (t's are spacer T nucleotides.). They were ordered from IDT with 5' fluorophore (Atto488, Atto565, Alexa647 or Alexa750), and 3' inverted dT modification, with HPLC purification. They are named as i.primerID#*. Sequences are listed in **Table S2**.

**Table S2. Imager strands used for fluorescent visualization.**

| Imager ID | Imager sequence |
|---|---|
| i.25* | /Fluorophore/tt-TATTATTGG-t -ATTATTGG-t /3InvdT/ |
| i.26* | /Fluorophore/tt-TAGGTTTAT-t-TAGGTTTAT-t /3InvdT/ |
| i.27* | /Fluorophore/ tt-ATGATGATG-t-ATGATGATG-t 3InvdT/ |
| i.28* | /Fluorophore/ tt-GTTAAGTTG-t-GTTAAGTTG-t/3InvdT/ |
| i.29* | /Fluorophore/ tt-GATTTTAGA-t-GATTTTAGA-t/3InvdT/ |
| i.30* | /Fluorophore/ tt-GAGAGTATT-t-GAGAGTATT-t/3InvdT/ |
| i.31* | /Fluorophore/ tt-AGTGAATAA-t-AGTGAATAA-t/3InvdT/ |
| i.32* | /Fluorophore/ tt-GAAAAAAAG-t-GAAAAAAAG-t/3InvdT/ |
| i.33* | /Fluorophore/ tt AATAGAAGGt AATAGAAGG-t /3InvdT/ |
| i.34* | /Fluorophore/ tt-GTAGTAGAG-t-GTAGTAGAG-t /3InvdT/ |
| i.35* | /Fluorophore/ tt-GAGTTTTTA-t-GAGTTTTTA-t /3InvdT/ |
| i.36* | /Fluorophore/ tt-AGATTAGTT-t-AGATTAGTT-t /3InvdT/ |
| i.37* | /Fluorophore/ tt-GAAGAGAAA-t-GAAGAGAAA-t /3InvdT/ |
| i.38* | /Fluorophore/ tt-TAGTATGTT-t-TAGTATGTT-t /3InvdT/ |
| i.39* | /Fluorophore/ tt-GTAAATGAA-t-GTAAATGAA-t /3InvdT/ |
| i.40* | /Fluorophore/ tt-TTGTAGGAT-t-TTGTAGGAT-t /3InvdT/ |
| i.41* | /Fluorophore/ tt-TTTTGATTG-t-TTTTGATTG-t /3InvdT/ |
| i.42* | /Fluorophore/ tt-GTTTGTAAG-t-GTTTGTAAG-t /3InvdT/ |
| i.43* | /Fluorophore/ tt-GTTATTTGT-t-GTTATTTGT-t /3InvdT/ |
| i.44* | /Fluorophore/ tt-GGTAGAAAA-t-GGTAGAAAA-t /3InvdT/ |
| i.45* | /Fluorophore/ tt-AAATAAGGG-t-AAATAAGGG-t /3InvdT/ |
| i.46* | /Fluorophore/ tt-AATGAAAGA-t-AATGAAAGA-t /3InvdT/ |
| i.47* | /Fluorophore/ tt-GAGTAAGAA-t-GAGTAAGAA-t /3InvdT/ |
| i.48* | /Fluorophore/ tt-GATTTATGG-t-GATTTATGG-t /3InvdT/ |
| i.49* | /Fluorophore/ tt-GGATAAATG-t-GGATAAATG t /3InvdT/ |
| i.50* | /Fluorophore/ tt-GTGAAGTAT-t-GTGAAGTAT t /3InvdT/ |
| i.51* | /Fluorophore/ tt-TTAGAGGTA-t-TTAGAGGTA-t /3InvdT/ |
| i.52* | /Fluorophore/ tt-AAATAGGAG-t-AAATAGGAG-t /3InvdT/ |

| | |
|---|---|
| i.53* | /Fluorophore/ tt-TTTGGATAG-t-TTGGATAG-t /3InvdT/ |
| i.54* | /Fluorophore/ tt-GATAGGGAT-t-GATAGGGAT-t /3InvdT/ |
| i.55* | /Fluorophore/ tt AAGTAATGA-t-AAGTAATGA t /3InvdT/ |
| i.56* | /Fluorophore/ tt GAGATTTAG-t-GAGATTTAG t /3InvdT/ |
| Test-imager | /Fluorophore/ tt-AATAAGAGA-t-AATAAGAGA-t /3InvdT/ |

***Branching primers.*** For stable hybridization of the secondary (branching) concatemers onto the primary concatemers, trimers of the unit repeat sequence were used as bridges, creating a 30-mer hybridization sequence. Hence, branching primers are designed in the format: 5'- p.1* - t -p.1* - t - p.1* - ttt (spacer) - p.2 - 3', where p.1 is the primer used for the primary concatemer, p.2 is the primer for the secondary concatemer, and t's are spacer T nucleotides. For secondary concatemers, we utilize extensions <500 bases based on empirical experience.

Similarly for iterative branching, the tertiary concatemer is designed to use the trimers of primer2 as the bridge, in the format: 5'- p.2* - t - p.2* - t  - p.2* - ttt - p.3 - 3', where p.3 is the primer for the tertiary concatemer and t's are spacer T nucleotides. For tertiary concatemers. For the third round, we utilize extensions <300 bases based on empirical experience.

### 2. Antibody-DNA conjugation and purification

***Conjugation.*** The conjugation involves crosslinking of thiol-modified DNA oligonucleotides to lysine residues on antibodies in a non-sequence-specific way. Briefly, 25 µl of 1 mM 5'-thiol-modified DNA oligonucleotides (Integrated DNA Technologies) were activated by 100 mM DTT (ThermoFisher #20291) for 2 h at room temperature in dark, and then purified using NAP5 columns (GE Healthcare Life Sciences #17-0853-02) to remove excessive DTT. Antibodies formulated in PBS only (or with sodium azide) were concentrated using 0.5 mL 50 kDa Ambicon Ultra Filters (EMD Millipore #UFC510096) to 2 mg/ml and reacted with maleimide-PEG2-succinimidyl ester crosslinkers (ThermoFisher #22102) for 2 h at 4°C (100 µg antibodies: 3.75 µl of 1 mg/ml crosslinker). Antibodies were then purified using 0.5 ml 7 kDA Zeba desalting columns (ThermoFisher #89883) to remove excessive crosslinkers. Activated DNA oligonucleotides were incubated with antibodies (11:1, DNA: Antibody molar ratio) overnight at 4°C. Final conjugated antibodies were washed using 2 ml 50 kDa Ambicon Ultra Filters six times to remove non-reacted DNA oligonucleotides. The list of bridge sequences used for conjugation is provided in **Table S3**. The list of antibodies and the corresponding bridge sequences used for each staining, as well as the capture and toehold strands for purification are provided in **Table S4**. Conjugated antibodies were diluted in the 1:50-1:200 for immunostaining.

***Information note regarding antibody-DNA conjugations.*** Not all commercial antibodies are provided in a formulation readily available for conjugation (for example antibodies may be provided in unpurified whole serum form or formulated with stabilizers or protectors that interfere with conjugation). Hence customized formulation of antibodies may be required. In addition, we currently utilize non-specific conjugation to Lys residues and provide a simple protocol to prepare custom conjugation of antibodies[6]. Although multiple DNA oligos can be attached to each antibody molecule for further signal amplification, our reaction conditions are optimized to achieve 1-2 oligos per antibody, to prioritize conserving the antigen recognition capability upon conjugation. Alternatively, site-specific conjugation chemistries could be utilized, including click labeling of antibody gylcosyl residues (available as the SiteClick™ kit from ThermoFisher). Independent of the conjugation method, we recommend testing of antibodies after conjugation to ensure functionality through comparison of the staining pattern with unconjugated antibody. As the high potential of DNA barcoding gains higher recognition and visibility, commercial antibody-DNA conjugation services and ready-to-use kits are also becoming available. Additionally, alternative recent probes (recombinant antibodies, nanobodies, aptamers, etc.) and probe labeling methods (such as unnatural amino acid incorporation or engineering of site-specific adaptor molecules) could facilitate new and highly-efficient means for standardized large-scale probe libraries as future resources.

***Purification.*** To increase the staining efficiency, conjugated antibodies can be optionally purified using a DNA toehold-mediated affinity pull-down protocol **(Fig. S1a)**. For this, 200 µl of high capacity streptavidin agarose (ThermoFisher #20357) was centrifuged down, washed 3 times using 500 µl PBS, and incubated with 10 µl 1 mM of biotin-labeled binding sequences in 300 µl PBS with 0.1% Triton X-100 for 30 min at room temperature. The agarose was then washed twice with PBS with 0.1% Triton X-100, followed by blocking with 250 µl blocking buffer (2% BSA + 0.1% Triton in PBS) for 1 h at room temperature with rotation. The agarose was then centrifuged and resuspended with 200 µl incubation buffer (1% BSA + 0.1% Triton in PBS) containing the DNA-conjugated antibodies, followed by rotation at 4°C for 1 h. The sample was centrifuged at 4°C and washed twice with 200 µl incubation buffer. The bound antibodies were recovered by adding 20 µl of 1 mM toehold strands (listed in **Table S3**) in 200 µl incubation buffer. After centrifugation, the supernatant was collected and the agarose was washed three times with 300 µl washing buffer (PBS + 0.1% Triton), collecting supernatant for each time. The supernatant was pooled together and buffer exchanged using 2 ml 50 kDa Amicon Ultra Filters six times to remove toehold DNA oligonucleotides. Binding sequences and toehold sequences were designed using NUPACK[7-10] and are provided in **Table S4**.

***Gel electrophoresis.*** To examine DNA antibody conjugation, antibodies were denatured in LDS sample buffer (ThermoFisher #NP0007) without reducing reagents (e.g. DTT or 2-ME) at 90 to 95°C for 3 min, and left to cool down to room temperature. The samples were run on 3 to 8% Tris-acetate PAGE gels (ThermoFisher #EA03752BOX) at 80 V for 30 min and 120 V for 3.5 h. The gels were stained with SimplyBlue™ safe stain (ThermoFisher #LC6060) according to the manufacturer's manual, and imaged using a Biorad Gel Doc™ EZ imager system. It should be noted that BSA should be avoided in the purification step if the sample needs to be examined using PAGE gels.

**Table S3. Bridge sequences used for antibody conjugation.**

| Bridge sequence index | DNA sequence |
|---|---|
| bc42_0 | AATTCTATGACACCGCCACGCCCTATATCCTCGCAATAACCC |
| bc42_1 | ATTATCCCTACCGCCAAATCTCCGTGTCCTTAACCGACCTAT |
| bc42_2 | CGTTATCGCCGCCTTATCCACTGTACGATCCTATTCCTCTCC |
| bc42_3 | GTTTCCTATATTTAGCGTCCGTGTCGTTCTCCCGCGCAACAG |
| bc42_4 | TATCTTAAGTCTTCGCGTGTTGTCTCGTCTGGGTATTGCGTT |
| bc42_5 | TCCTGTCCCGACGATCCTACCCTTAAAGTTACTGCGCACCCT |
| bc42_6 | CGGTGAGGTAGGAGTCGTGCGTATCGTTTCCTATATAGCCGT |
| bc42_7 | AGTTCCTGTAGTATCCCGTCGCATAGTCGTACATTCACCGTC |
| bc42_8 | AACAATTCAGCTCCGCCTTATACCGTCTTACCGCCAACATCG |
| bc42_9 | GAATTTGGCCCGTTCTATGTCTAACTCGTGTTCCCGCTTGTA |
| bc42_10 | GTCCTCGCTCTTTCCGCATTTTCCCGTATGCGCTTTGTATTA |
| bc42_11 | TGTCTAAATTCTAATGCCGCCCTATGCCGCCGTTCCAACAAT |
| bc42_12 | CCTTCCGCCGTATGAATTTGACCCGAAGCCCAACCCGACCCT |
| bc42_13 | CAGTTCTTGTATCGCGTCACTTATCGGTTATTGTCCTCTCGC |
| bc42_14 | CCAACCTCTCGTACCAAATTCCGCCACTCAAGCCGTATCAAA |
| bc42_15 | GTTTCAAGAGTCCGTCGCAAATTCCACTACACGCTACGCCCA |
| 25mer-tester | TATTTAGTGTTCGAATAGTTCGATCTAG |

The full set of 84 designed bridge sequences are available in our recent work[4]. All sequences were ordered from IDT.

**Table S4. Antibodies used in SABER experiments, conjugated bridge sequences and respective SABER primers.**

| Antibody target | Source | Bridge sequence (for conjugation) | Capture and Toehold sequences for purification (if purified) | SABER primer sequences used in the experiments (+ denotes branching) |
|---|---|---|---|---|
| Cone arrestin | Millipore #AB15282 | 25mer-tester | Capture: Biotin-GTTGCTGTCGTATGT-CTAGATCGAACTATTC | p.30 (Fig. 2) or p28 + p25 (Fig. 3) or test-primer |

| | | | | |
|---|---|---|---|---|
| | | | Toehold: GAATAGTTCGATCTAG-ACATACGACAGCAAC | (Fig. 6) |
| GFAP | ThermoFisher #13-0300 | bc42_1 | Capture: Biotin - GGGTAGGGTAGTGGT-ATAGGTCGGTTAAGGA Toehold: TCCTTAACCGACCTAT-ACCACTACCCTACCC | p.36 (Fig. 6 and Fig. 7) |
| PKCα | Novus #NB600-201 | bc42_2 | Binding sequence: Biotin - CGAGTGAGGTGGAAT-GGAGAGGAATAGGATC Toehold sequence: GATCCTATTCCTCTCC-ATTCCACCTCACTCG | p.25 (Fig. 6) |
| SV2 | HybridomaBank | bc42_3 | Capture: Biotin - CGAGTGGTAAGGCAT-CTGTTGCGCGGGAGAA Toehold: TTCTCCCGCGCAACAG-ATGCCTTACCACTCG | p.26 (Fig. 6, Fig.S6, Fig. 7 and Fig. S7); or p27 + p28 + p32 (Fig. S3) |
| Collagen IV | Novus #NB120-6586 | bc42_4 | Capture: Biotin - GAAATAGAATGAACG-AACGCAATACCCAGAC Toehold: GTCTGGGTATTGCGTT-CGTTCATTCTATTTC | p.27 (Fig. 6 and Fig.7) |
| Rhodopsin | EnCor Bio #MCA-A531 | bc42_7 | Capture: Biotin - GTTAAGGTGGAATGA-GACGGTGAATGTACGA Toehold: TCGTACATTCACCGTC-TCATTCCACCTTAAC | p.33 (Fig. 6 and Fig. 7) |
| Calbindin | EnCor Bio #MCA-5A9 | bc42_8 | Capture: Biotin - GGTGAGGTGTAGTGG-CGATGTTGGCGGTAAG Toehold: CTTACCGCCAACATCG-CCACTACACCTCACC | p.34 (Fig. 6) |
| Vimentin | Cell Signaling #5741S | bc42_9 | Capture: Biotin - CGGAACAGATAAAGA-TACAAGCGGGAACACG Toehold: CGTGTTCCCGCTTGTA-TCTTTATCTGTTCCG | p.28 (Fig. 6 and Fig. 7) |
| Calretinin | EnCor Bio #MCA3G9 | bc42_10 | Capture: Biotin - GCCAAATTCCACCGC-TAATACAAAGCGCATA Toehold: TATGCGCTTTGTATTA-GCGGTGGAATTTGGC | p.30 (Fig. 6 and Fig. 7) |
| VLP1 | EnCor Bio #MCA-2D11 | bc42_11 | Capture: Biotin - CGGATGATGAGGGTG-ATTGTTGGAACGGCGG Toehold: CCGCCGTTCCAACAA-TCACCCTCATCATCCG | p.39 (Fig. 6) |
| Alpha-Tubulin | ThermoFisher #MA1-80017 | bc42_0 | Capture: Biotin - GTTGAGTGAGGTTGA-GGGTTATTGCGAGGAT Toehold: ATCCTCGCAATAACCC-TCAACCTCACTCAAC | p.30 |
| Ki-67 | Cell Signaling #9129 (formulated in PBS) | bc42_1 | Capture: Biotin - GGGTAGGGTAGTGGT-ATAGGTCGGTTAAGGA Toehold: TCCTTAACCGACCTAT-ACCACTACCCTACCC | p.30 + p.28 + p.25 (Fig. 3e-j, S3h), or p.41 + p.34 (Fig. 5c), or p.30 or p.30 + p.28 (all other figures) |

| | | | | |
|---|---|---|---|---|
| CD8a | Cell Signaling #85336 (formulated in PBS) | bc42_2 | Capture: Biotin - CGAGTGAGGTGGAAT-GGAGAGGAATAGGATC Toehold: GATCCTATTCCTCTCC-ATTCCACCTCACTCG | p.40 + p.28 (Fig. 5c) or p.25 + p.31 (all other figures) |
| PD-1 | Cell Signaling #43248 (formulated in PBS) | bc42_3 | Capture: Biotin - CGAGTGGTAAGGCAT-CTGTTGCGCGGGAGAA Toehold: TTCTCCCGCGCAACAG-ATGCCTTACCACTCG | p.26 + p.39 |
| IgA | Jackson #109-005-011 | bc42_7 | Unpurified | p.34 (Fig. 5a) or p.25 (Fig. 5c) |
| CD3e | Cell Signaling #85061 (formulated in PBS) | bc42_9 | Capture: Biotin - CGGAACAGATAAAGA-TACAAGCGGGAACACG Toehold: CGTGTTCCCGCTTGTA-TCTTTATCTGTTCCG | p.27 + p.32 |
| IgM | Jackson #709-006-073 | bc42_11 | Unpurified | p.39 (Fig. 5a) or p.35 (Fig. 5c) |
| Bassoon | Enzo ADI-VAM-#PS003 | bc42_0 (conjugated onto the secondary antibody, Jackson ImmunoResearch #715-005-151) | Unpurified | p.30 (Fig.7 and Fig. S7) |
| Homer1b/c | ThermoFisher #PA5-21487 | bc42_3 (conjugated onto the secondary antibody, Jackson ImmunoResearch #711-005-052 ) | Unpurified | p.26 (Fig.7 and Fig. S7) |
| Anti-rabbit IgG (to detect Ki-67 indirectly) | Jackson # 711-005-152 | bc42_3 | Unpurified | p.30 + p.28 (Fig. 3h-j, S3i) |

Antibodies used to validate colocalization of VLP1 and Calretinin in Fig. **S6d-f**: Calretinin (SantaCruz #SC-365956; EnCor Bio #CPCA-Calret; EnCor Bio #MCA-3G9 AP), VLP1 (EnCor Bio #RPCA-VLP1; EnCor Bio #CPCA-VLP1; EnCor Bio #MCA-2D11).

Fluorophore-conjugated secondary antibodies used for reference imaging: anti-rat-Alexa647 (ThermoFisher #A-21472), anti-rabbit-Alexa488 (ThermoFisher #A-21206), anti-rabbit-Atto488 (Rockland #611-152-122S), anti-mouse-Alexa647 (ThermoFisher #A-31571).

### 3. Microtubule staining in cell culture and FWHM analysis

***Staining.*** BS-C-1 cells were plated on eight-well ibidi glass-bottom μ-slides (ibidi #80826) and grown until 50-60% confluency. For cell culture experiments in **Fig. 2** and **Fig. 3**, cells were fixed with 4% paraformaldehyde (PFA) for 45 min, and quenched with 100 mM $NH_4Cl$ in PBS for 20 min and washed with PBS for 5 min. Cells were then permeabilized and blocked in 2% nuclease-free BSA (AmericanBIO, CAS 9048-46-8) + 0.1%-Triton in PBS for 30 min. Samples were incubated with DNA-conjugated primary antibodies diluted in the incubation buffer made of 0.1% Triton X-100, 2% nuclease-free BSA, 0.2 mg/ml sheared salmon sperm DNA, 0.05% Dextran sulfate (Millipore #S4030), 4 mM EDTA in PBS overnight at 4°C, and then washed with PBS with 0.1% Triton X-100 and 2% BSA for 3 times for 10 min each. Samples were then washed with PBS twice for 5 min and post-fixed using 5 mM BS(PEG)$_5$ (ThermoFisher #21581) in PBS for 30 min, followed by quenching in 100 mM $NH_4Cl$ in PBS for 5 minutes. The incubation with the primary concatemer was performed at 37°C in 20% formamide (deionized, Ambion #AM9342), 10% Dextran sulfate (Millipore #S4030) and 0.1% (v/v) Tween-20 in 2× SSC with 0.2 mg/ml sheared salmon sperm DNA for 3 h. 650 base long primary concatemers prepared *in vitro* by PER were diluted in this buffer at 133 nM final primer concentration (primer concentration in the PER mix is considered a proxy for the concatemer concentration after the reaction, since all primers are expected to be extended by the catalytic hairpins that are provided in excess). After concatemer hybridization the samples were washed for 5 min at RT with 45% formamide in PBS and three times for 10 min each with PBS + 0.1% Triton X-100 at 37°C. Branching hybridization was performed at 37°C in 30% formamide, 10% Dextran sulfate and 0.1% (v/v) Tween-20 in 2× SSC with 0.2 mg/ml sheared salmon sperm DNA for overnight at 133 nM final concentration of the 450 base long secondary concatemers. Samples were washed for 5 min at room temperature with 45% formamide in PBS and three times for 10 minutes each with PBS + 0.1% Triton X-100 at 37°C. Imagers were hybridized at 1-1.5 µM final concentration in PBS + 0.1% Triton X-100 for 1 h at room temperature (hybridization duration with the imagers can be significantly decreased for faster preparation), followed by a 5 min wash with PBS + 0.1% Triton X-100 and two times 5 min wash with PBS. Samples were stained with 1 µg/ml DAPI (Invitrogen #D1306) in PBS for 10 min and washed twice for 1 min with PBS. Imaging was performed in SlowFade with DAPI (Invitrogen #S36938) embedding medium.

***Imaging.*** For the images exemplified in **Fig. 2a**, **S2a** and **S3a**, a Leica SP5 confocal with 63×/NA 1.3 Glycerol objective was used. A white light laser (470-670 nm) was used at 650 nm for excitation of Alexa647, and a PMT was used for detection.

***FWHM Analysis.*** The best focus planes for isolated microtubules were manually selected from z-stacks. 2.5 µm lineplots were drawn across isolated microtubules in this single plane image using Fiji[3]. Also small rectangular background areas were manually drawn to obtain the background values around selected microtubule lines. Average background was subtracted from the lineplot values. Gaussian curves were fit to the background-subtracted values and FWHM was calculated based on the fits and average values were obtained by analyzing 30-45 lineplots from 5 images per condition using a Python script. A similar calculation was performed for 200 nm dark red fluosphere beads (Invitrogen #F8807). Distributions were displayed as box plots. Lineplots with multiple discernible peaks were discarded. Two-sample t-test was performed to check for statistical significance.

### 4. Preparation, staining, SABER application, quantification and analysis of human tonsil FFPE sections

***Preparation of formalin-fixed paraffin-embedded (FFPE) tonsil samples and antigen retrieval.*** Human specimens were retrieved from the archives of the Pathology Department of Beth Israel Deaconess Medical Center under the discarded/excess tissue protocol as approved in Institutional Review Board (IRB) Protocol #2017P000585. Five micron sections were cut with a rotary microtome, collected in a water bath at 30˚C, transferred to positively charged glass slides and baked at 60˚C for 2 h. For antigen unmasking, slides were placed on a PT-Link instrument (Agilent), which allows the entire pre-treatment process of deparaffinization, rehydration and epitope retrieval (with citrate buffer) to be combined into a single step. Slides were held at 4˚C in PBS until staining. Antigen-retrieved FFPE sections used in multicolor experiments in **Fig. 5** were acquired from Ultivue Inc.

***Staining of antigen-retrieved FFPE tonsil samples.*** After antigen retrieval, sections were optionally stored in PBS at 4˚C up to 2 weeks. For staining, sections were washed in PBS for 15 min and outlined with a hydrophobic pen (ImmEdge Hydrophobic Barrier PAP Pen, Vector Laboratories #H4000) or be enclosed in a removable chamber (ibidi, #80381). At this stage a mild 1 h pre-bleaching with 1% $H_2O_2$ in PBS can be optionally applied. This step may help to reduce the overall sample autofluorescence, but is not necessary for Immuno-SABER. We incorporated this step for the Ki-67 stainings in **Fig. 3** and **S3**, to keep the conditions similar to TSA preparations, which benefit from a pre-bleaching step significantly. Samples were blocked for 1 h with PBS containing 2% BSA and 0.1-0.3% Triton X-100, with 3 buffer exchanges. DNA-conjugated primary antibodies were diluted in the blocking solution supplemented with 0.2 µg/ml sheared salmon sperm DNA, 0.05% Dextran Sulfate, and optionally 0.4 mM EDTA, and incubated on the samples overnight at 4˚C in a humidified chamber. The antibodies and respective bridge sequences used for the experiments are listed in **Table S4**. Depending on the antibody this step can be shortened to 1 h when performed at room temperature or at 37°C. Excess antibodies were washed at room temperature 3 times for 15 min each with PBS containing 2% BSA and 0.1-0.3% Triton X-100, and 2 times for 5 min with PBS. Bound antibodies were then post-fixed with 5 mM BS(PEG)$_5$ in PBS for 30 min at RT, followed by quenching in 100 mM $NH_4Cl$ in PBS for 5 minutes and washed for 15 min with PBS with 0.1% Triton X-100 at RT. Post-fixation is critical to ensure that the antibodies are not washed during further labeling and imaging. The incubation with the primary concatemer was performed at 37°C in 20-30% formamide, 10% Dextran sulfate and 0.1% (v/v) Tween-20 in 2× SSC with 0.2 mg/ml sheared salmon sperm DNA for 1 h to overnight. Concatemers prepared *in vitro* by PER were diluted in this buffer at 66-150 nM (1:15 to 1:7.5 dilution) final primer concentration. For multiplexing, all primary concatemers were incubated simultaneously. After concatemer hybridization the samples were washed for 5 minutes at room temperature with 45-50% formamide in PBS and three times for 10 minutes each with PBS + 0.1% Triton X-100 at 37°C. For cases, where further amplification was desired, branching hybridizations were performed similarly (at 37°C in 30% formamide, 10% Dextran sulfate and 0.1% Tween-20 in 2× SSC with 0.2 mg/ml sheared salmon sperm DNA for 1 h to overnight). Bridge and primer sequences for each target and experiment are given in **Table S4**.

For the unamplified sample, the unextended primer with one imager binding site (equivalent to two repeats of the primer sequence) was incubated at the same concentration instead of the extended concatemer.
For CD8a, bc42_2*-tt-p.25-a-p.25-a:
5'-GGAGAGGAATAGGATCGTACAGTGGATAAGGCGGCGATAACG-tt-CCAATAATA-a-CCAATAATA a-3'
For Ki-67, bc42_0*-tt-p.30-a-p.30-a:
5'-GGGTTATTGCGAGGATATAGGGCGTGGCGGTGTCATAGAATT-tt-AATACTCTC-a-AATACTCTC a-3'

For different tissue types, and combination of targets for multiplexing, experimental conditions may need to be optimized to achieve the best signal level. For primary concatemers we recommend using sequences ≤650 bases, for secondary concatemers ≤450 bases, and for tertiary concatemers ≤250 bases. For iterative amplification the wash temperatures were raised to 42°C and the 45-50% formamide in PBS wash step was performed once as a final wash only at the end of all iterations (rather than after each amplification round).

***Fluorophore hybridization and dehybridization.*** Imagers were hybridized at 1-1.5 μM final concentration in PBS + 0.1% Triton X-100 for 1-2 h at RT, followed by a 5 min wash with PBS + 0.1% Triton X-100 at 37°C and two times 5 min wash with PBS at room temperature. Elongated hybridization times were used for convenience, for faster protocols, hybridization duration for the imagers can be significantly decreased with similar performance. Samples were stained with 1 μg/ml DAPI in PBS for 10 min and washed twice with PBS for 1 min each. For multiplexing experiments samples were imaged in PBS shortly after preparation and coverslips were temporarily secured with Fixogum (Marabu). Coverslipping of the samples for imaging is optional depending on the instrument type. Imagers were removed with 10 min incubation at room temperature in 50% formamide in PBS, followed by two times 5 min wash with PBS at room temperature. A new round of imager hybridization was performed as above.

For experiments with a single round of imager hybridization (no multiplexing or only spectral multiplexing) samples were embedded in SlowFade with DAPI, secured with nail polish, and imaged on the same day or embedded with ProLong Diamond (Invitrogen #P36971) and incubated at RT overnight for curing.

***Tyramide signal amplification.*** For FFPE samples, Anti-Rabbit IgG Alexa647 Tyramide SuperBoost™ Kit (Life Technologies #B40916) was used according to manufacturer's recommendations. The optional step of 1 h pre-bleaching with 1% $H_2O_2$ in PBS was applied. Tyramide-Alexa647 was incubated for 2.5 min, 7.5 min and 10 min. Longer incubation times (≥7.5 min) were observed to cause increased blurring of the signal, making shorter incubation times more favorable (as suggested by the manufacturer).

***Fluorescence Imaging.*** Quantification samples in **Fig. 2**, **3**, **S2** and **S3, S5a** were imaged with an Olympus VS-120 system equipped with a Orca R2 monochrome (16 bit) camera using a 20×/0.75 NA air objective, with a pixel size of 0.320 μm and single plane tile scans were acquired. For Fig. 3 stainings, exposure times of 35 ms and 30 ms were used for DAPI and Ki-67 respectively. For the CD8a stainings in **Fig. 1-2**, exposure time of 350 ms was used.

For the zoom-in's in **Fig. 3j** and **S3h** a Leica SP5 confocal with 63×/NA 1.3 Glycerol objective were used. A white light laser (470-670 nm) was used at 650 nm for excitation of Alexa647, and a PMT was used for detection.

Multiplex FFPE samples in **Fig. 5** (and **S5b**) were imaged with a Perkin Elmer Vectra Polaris microscope in whole slide scanning configuration with a 20×/0.80 NA air objective. For **Fig. 5c** imagers were first hybridized for three of the targets (as depicted in the figure), followed by nuclear DAPI staining, washing, coverslipping and whole section imaging in 5 channels (3 markers plus DAPI and autofluorescence). This was followed by dehybridization as described above. The second set of imagers was similarly hybridized and the sample was reimaged.

Images obtained in .qptiff format were converted to .tif with Imaris software and aligned using "Align by line ROI" plugin on Fiji using the DAPI fluorescence in each cycle and overlays for display were assembled in Adobe Photoshop.

***Signal quantification.*** For quantification of signal amplification for CD8a labeling in tonsil samples in the **Fig. 2** and **3**, rectangular regions of interest (ROIs) covering 0.30-1.2 $mm^2$ tile scans were selected after manual inspection to exclude areas with autofocusing errors or sectioning imperfections, and a customized CellProfiler routine was used first to remove masked autofluroescent structures identified in an independent channel (using global robust thresholding), then to calculate mean fluorescence signal/pixel for cell regions masked via thresholding of the CD8a signal[11]. For masking of the labeled structures, edges were enhanced by Canny edge finding method with automatic threshold calculation. Then global robust background thresholding was applied. Upper outlier fraction and correction factor for thresholding was adjusted manually to compensate for the difference in overall signal level under different amplification conditions.

The background was calculated by averaging the CD8a fluorescence signal of the tissue regions (ensured by presence of DAPI) outside of the masked signal regions (after dilation of the masked pixels). The final intensity value was obtained by subtracting the mean background value for each ROI from the average signal value for masked cells. Amplification fold was calculated by dividing the mean background-subtracted fluorescence by the unamplified sample (for linear SABER, **Fig. 2c,e**) or by the linear amplification sample (for branched SABER, **Fig. 3b,d**).

Figures were prepared for display using OMERO for ROI selection, and scaling[12].

***Nuclear segmentation and analysis of single-nuclei intensity distribution and quantification of fold-amplification.*** We annotated the contours of all the nuclei for 80 patches of 128×128 pixels. This dataset was then augmented 8-fold using reflections and rotations, resulting on a dataset of 640 images. For each image, an equal number of non-contour pixels were randomly selected to represent the complementary class in the machine learning algorithm. We trained a variant of the U-Net model[13], adding batch normalization[14] and residual learning[15], to classify pixels in two classes (class 1: nuclei contours; class 2: background and nuclei interiors). We used our own implementation in TensorFlow with the following hyperparameters: Number of feature maps in first convolutional layer: 8; Number of feature maps in subsequent downsampling layers: 2 times the number of feature maps in previous layer; Downsampling and upsampling factor: 2; Convolution kernel size: 3; Number of extra convolutions in each layer: 1; Variance of truncated normal distribution generating initial random weights: 0.1; Number of downsampling layers: 2; Batch size: 8; Number of training steps: 20000; Learning rate: initially 0.1, with 'staircase' exponential decay (step 1000, rate 0.95), and momentum 0.9.

The approximate center of each nucleus was then identified through the regional maxima of a Gaussian-blurred version of the inverted contour probability class. These regional maxima were used as seed points for applying a marker-controlled watershed transform. Background objects were eliminated based on mean intensity and area. The mean, median, and 95th percentile fluorescence intensity in each segmented nucleus were measured and exported in .csv format. For the fold amplification estimation, mean signal intensity from all the nuclei in each tissue sample were summed up to obtain the total Ki-67 signal level. Then, total signal for each sample was divided by the total signal for the unamplified condition. Total values were not corrected for the variation of the number of cells in each section (tissue samples are consecutive sections from the same tissue). Histograms that show the distribution of Ki-67 signal per individual cells were plotted using Python.

**5. Preparation, staining, SABER application and quantification on mouse retina cryosections**

***Sample preparation.*** All animal procedures were in accordance with the National Institute for Laboratory Animal Research Guide for the Care and Use of Laboratory Animals and approved by the Harvard Medical School Committee on Animal Care. Animals were given a lethal dose of sodium pentobarbital (120 mg/kg) (MWI, 710101) and enucleated immediately. Eyes were removed and fixed in PFA for 15 to 30 min. Following dissection, retinas were immersed in 30% sucrose overnight prior to freezing in TFM (EMS, 72592) and cryosectioning at ~30-40 µm. Eight-well ibidi glass-bottom µ-slides were treated with 0.3 mg/ml poly-D-Lysine for at least 30 minutes, followed by 3 times PBS washes. Retina sections were immobilized onto the glass and stored at -20°C. Sections were washed with Tris-buffered saline (TBS) + 0.3% Triton X-100 for three times with 10 min per wash. They were then blocked and stained as above, except that BSA was replaced by 5% donkey serum and 0.1% Tween-20 was replaced by additional 0.2% Triton X-100.

***Immuno-SABER application.*** PER extensions were diluted in 1:7.5 to 1:20 (depending on the target density) in incubation buffers. Two different concatemer hybridization buffers were used: buffer 1 is 40% formamide + 10% Dextran sulfate + 0.1% Triton X-100 + 0.02% sodium azide + 5mM EDTA in PBS, and buffer 2 is 30% formamide + 10% Dextran sulfate + 0.1% Triton X-100 + 0.02% sodium azide + 5 mM EDTA in PBS. Buffer 1 was used for inucbation with primary concatemers and buffer 2 was used for branching concatemers. The samples were left at room temperature overnight in a humidified chamber. The samples were washed with 45% formamide + 0.1 % Triton X-100 + 5mM EDTA in PBS for 30 minutes and twice with 30% formamide + 0.1 % Triton X-100 + 5mM EDTA in PBS for 30 min at room temperature. For branched conditions, 45% formamide was replaced with 40% formamide for the first wash. For multiplexed imaging experiment, all primary concatemers were incubated simultaneously. Bridge and primer sequences for each target and experiment are given in **Table S4**.

To quantify linear SABER amplification for cone arrestin, the amplification samples were incubated with SABER concatemers extended from 25mer-tester*-tt-p.28 (CTAGATCGAACTATTCGAACACTAAATA-tt-CAACTTAAC). The unamplified samples were hybridized with the unextended primer 25mer-tester*-tt-p.28-a-p.28 (CTAGATCGAACTATTCGAACACTAAATA-tt-CAACTTAAC-a-CAACTTAAC), which carries one imager binding site (equivalent to two repeats of the primer sequence) instead of the extended concatemer (at the same final concentration).
For quantification of branched SABER for cone arrestin, the branching sample was first incubated with the primary concatemers extended from 25mer-tester*-tt-p.28, followed by incubation with secondary concatemers extended from 28*-t-28*-t-28*-ttt-p.25 primer (GTTAAGTTG-t-GTTAAGTTG-t-GTTAAGTTG-ttt-CCAATAATA).

For iterative SABER quantification for SV2, the unamplified sample was hybridized with unextended primer bc42_3*-tt-p.27-a-p.27 (CTGTTGCGCGGGAGAACGACACGGACGCTAAATATAGGAAAC-tt-CATCATCAT-a-CATCATCAT); the linear amplified sample was hybridized with the primary concatemer extended from bc42_3*-tt-p.27 (CTGTTGCGCGGGAGAACGACACGGACGCTAAATATAGGAAAC-tt-CATCATCAT). The branching sample additionally hybridized with the secondary concatemer 27*-t-27*-t-27*-ttt-p.28 (ATGATGATG-t-ATGATGATG-t-ATGATGATG-ttt-CAACTTAAC); the iterative amplification sample was additionally hybridized the tertiary concatemer extended from 28*-t-28*-t-28*-ttt-p.32 (GTTAAGTTG-t-GTTAAGTTG-t-GTTAAGTTG-ttt-CTTTTTTTC).

***Fluorescence Imaging.*** Fluorophore-labeled imager strands were diluted in 0.1% Triton X-100 in PBS to ~250 nM-1 µM, and incubated with samples for 30 min, followed by washing using 0.1% Triton X-100 in 0.5× PBS for three times. Samples were left in PBS during image acquisition.

For 10-color multiplexing, the entire experiment was done using five rounds of buffer exchange with an average of 1.5 h per round that included 30 min for imager hybridization, 15 min for excessive imager strands removal, 15 min for imaging and 30 min for imager strands removal. It should be noted that the duration for each step is sample dependent with thicker samples requiring longer time to ensure complete penetration and signal removal. To ensure the best signal and simplify the multiplexed experimental design, we allowed excessive time for each step, however it is possible to shorten incubation and washing times upon optimization.

All images for mouse retina sections were acquired using a Zeiss Axio Observer with LSM 710 scanning confocal system with a 20×/0.8 NA air objective. The images were 512×512 pixels or 1024×1024 pixels and acquired at acquisition speed 7. Each image was acquired by averaging 2 images. Atto488 was visualized using a 488 nm laser; Atto565 was visualized using a 546 nm laser; Alexa647 was visualized using a 633 nm laser. To remove imager strands, samples were washed three times with 30% formamide + 0.1% Triton X-100 in 0.1× PBS. The samples were left in PBS during imaging. Acquired images were scaled and colorized for display using FIJI[3] and Photoshop.

***Tyramide signal amplification.*** For retina cryosections, Alexa Fluor 647 TSA™ Kit #6 with HRP–goat anti-mouse IgG (Life Technologies #T20916) was used following manufacturer's recommendations, without the optional pre-bleaching step and with 7.5 min tyramide incubation.

***Quantification.*** For quantification of signal amplification in retina samples (SV2 and cone arrestin), the mask regions were selected manually and mean fluorescence intensity was calculated. The background was calculated by averaging the fluorescence signal of six randomly selected regions outside the retinas. The final fold-amplification values were obtained by subtracting the background value from the average signal value for the condition and normalizing that by the unamplified (**Fig. 2e**, **S3e**) or by the linear condition (**Fig. 3c-d**).

**6. Preparation, staining, SABER application on whole mount retina samples**

***Sample preparation and staining.*** Whole mount retina samples were prepared as free-floating samples, which allowed reagents to penetrate from both sides of retinas. The thickness of whole mount retina samples are typically ~160 to 180 μm. The samples preparation was conducted with a similar protocol as above but with longer incubation and wash times. DNA-conjugated primary antibodies were incubated for 40 h at 4°C, and washed for 3 × 1 h. DNA extensions were incubated for 40 h at room temperature, and washed for 3 × 1 h. Fluorescent oligos were incubated for 2 h at room temperature, followed by 3 × 30 min. Retinas were flattened by creating 4 incisions and mounted on a glass slide. Bridge and primer sequences for each target are given in **Table S4**.

***Imaging.*** The images were acquired using a Zeiss Axio Observer with LSM 710 scanning confocal system with a 10×/0.45 NA air objective. A z-stack of 98 sections with 1 μm spacing was taken for each target. It should be noted that although the entire whole mount retina is about ~180 μm, the signal from vimentin and collagen IV are typically located in half of the retina section from the nerve fiber layer to the outer plexiform layer.

**7. PER sequence crosstalk analysis**

For microtubule stainings in **Fig. 4** and **Fig. S4**, BS-C-1 cells were grown in glassbottom 96-well plate (Ibidi #89626) with 5,000 cells per well. Cells were fixed with 4% PFA for 15 minutes, and quenched with 50 mM $NH_4Cl$ in PBS for 7 min. Cells were then permeabilized and blocked in 0.1% Triton X-100, 0.1% Tween20, 2% nuclease-free BSA (AmericanBIO, CAS 9048-46-8) and 0.2 mg/ml sheared salmon sperm DNA (ThermoFisher #AM9680) in PBS for 1 h. Samples were incubated with DNA-conjugated primary antibodies diluted in incubation buffer (0.05% Triton X-100, 0.05% Tween20, 2% nuclease-free BSA, 0.2 mg/ml sheared salmon sperm DNA, 0.05% Dextran sulfate (Millipore #S4030), 5 mM EDTA in PBS) overnight at 4°C, and then washed with washing buffer (0.05% Triton X-100, 0.05% Tween-20, 2% nuclease-free BSA, 5 mM EDTA in PBS) for five times (1-2 min for the first two washes and 10 min incubation for the other three washes). Samples were washed with PBS twice and post-fixed using 5 mM $BS(PEG)_5$ (ThermoFisher #21581) in PBS for 1 h, followed by quenching in TBS for 10 min.

For cognate wells, 20 nM corresponding imager strands were incubated, while for crosstalk wells, all other imager strands were incubated with 20 nM for each imager strand. The samples were imaged using Zeiss Axio Observer

Z1 with a 20×/0.8 air objective. For signal quantification, the bright field images were acquired and used to create masks in MATLAB. Average fluorescence signals were calculated within the mask region. The background was calculated as the average fluorescence signals outside the cells. The final fluorescence intensity was average fluorescence intensity within cell masks minus background intensity.

**8. Expansion microscopy**

The PER primer sequences were modified with acrydite at the 5'-end (Integrated DNA Technologies), and extended as above. Mouse retina samples were stained with DNA-conjugated antibodies, followed by SABER extension incubation. After washing away excessive SABER extension, a layer of expandable gel was formed according to the original expansion microscopy protocol[16]. In brief, samples were incubated monomer solution (1x PBS, 2 M NaCl, 8.625% (w/v) sodium acrylate, 2.5% (w/v) acrylamide, 0.15% (w/v) N,N'-methylenebisacrylamide) with ammonium persulfate (APS) and teramethylethylenediamine (TEMED) on ice with open air for 20 minutes. A gelation chamber was then constructed by placing a No.1 coverglass on each side of the tissue section and covering with a No.1 coverglass. The specimens were transferred to a humidified incubator and left at 37°C for 2 h. The samples were then digested using Proteinase K (New England BioLabs, Cat.No: P8107S) at 1:100 dilution in digestion buffer (50 mM Tris (pH 8), 1 mM EDTA, 0.5% Triton X-100, 0.8 M guanidine HCl) at 37 °C overnight. The digested samples were then expanded by placing in excess volumes of de-ionized water. To prevent expanded samples from shrinkage, they were re-embeded in a nonexpandable gel (3% acrylamide, 0.15% N,N′-Methylenebisacrylamide with 0.05% APS, 0.05% TEMED). The gel was placed on a bind-silane treated No.1.5 coverglass and immersed in the gel solution on ice for 20 minutes, followed by gelation at 37°C for 1.5 h. To coat coverglass with bind-silane, wash the coverglass with $ddH_20$ followed by 100% ethanol. Dilute 5 µl of Bind-Silane reagent (GE #GE17-1330-01) into 8 ml of ethanol, 1.8 ml of $ddH_2O$ and 200 µl of acetic acid. After re-embedding, the samples were imaged as above in regular retina tissue section imaging.

For the primary neuron culture, the culture was grown on a 12 mm diameter round #1 coverslips, and stained with Bassoon and Homer1b/c antibodies, followed by DNA-conjugated anti-mouse and rabbit secondary antibodies. The expandable gel (19% (w/v) sodium acrylate, 10% (w/v) acrylamide, 0.05% (w/v) N,N'-methylenebisacrylamide in PBS) was formed by placing the coverslip against a parafilm sheet with 20 µl of expansion gel solution in between. The gel was then digested and expanded as above. The gel was transferred to a coverslip dish (ibidi #81148) and was incubated with the imager strands (Atto488-i.30* and Atto565-i.26*) in 0.5× PBS without re-embedding and left in 0.5× PBS during confocal imaging, which gave a similar expansion factor of ~3 fold.
